# *Photorhabdus* metabolites reshape soil microbial communities and promote plant growth and insect resistance

**DOI:** 10.64898/2026.03.12.710065

**Authors:** Jaspher Ewany, Ivan Hiltpold, Emmanuel Defossez, Gaétan Glauser, Carla C. M. Arce, Wei Zhang, Sergio Rasmann, Ted C.J. Turlings, Ricardo A.R. Machado

## Abstract

*Photorhabdus* bacteria are potent insect-killing microbes associated with entomopathogenic nematodes and offer opportunities for environmentally benign pest control. They can be applied as foliar sprays or soil drenches without their nematode vector, resulting in massive amounts of *Photorhabdus* cells and their (toxic) metabolites introduced into the soil. However, their effects beyond the target organisms are unknown. To fill this knowledge gap, we investigated the soil legacy effects of *Photorhabdus* cells and their metabolites on soil microbial communities, plant performance and resistance to herbivores. To this end, we first conditioned soils with i) mechanically killed (MK) or *Photorhabdus*-infected insect larvae, ii) aqueous extracts of MK or *Photorhabdus*-infected insect larvae, iii) cell-free *Photorhabdus* supernatants, iv) autoclaved soil complemented with live soil previously conditioned with MK or *Photorhabdus*-infected insect larvae. We then grew maize plants in these soils and measured plant biomass, profiled soil microbial communities and plant metabolites, and evaluated plant resistance against two pest insects *Diabrotica balteata* and *Spodoptera frugiperda*. We found that conditioned soils increased plant biomass by 10–26% relative to controls and significantly altered soil bacterial and nematode communities, and to a lesser extent, fungal communities. Re-inoculating conditioned soil microbiota into autoclaved soils recreated the plant growth-promoting effects, indicating microbial-mediated mechanisms. Additionally, plants grown in soils conditioned with *Photorhabdus*-infected insect cadavers were often more resistant to herbivorous insect attack, in a strain-specific manner. On average, *D. balteata* and *S. frugiperda* larvae gained 10–20% and 10–59% less weight, respectively, when fed on plants grown in conditioned soils than on plants grown in control soils. The plant metabolic profiles of plant leaves and roots also varied with resistance levels. We conclude that *Photorhabdus* metabolites modulate soil microbial communities towards a structure that enhances plant growth and triggers systemic responses against herbivores.

## Introduction

Insect pests are a significant constraint to global food production, causing 20 – 40% annual crop losses [1]. From a financial perspective, it is estimated that pests cause about $70 billion loss and that the total food consumed by pests can feed more than 1 billion people [1,2].

The fall armyworm (FAW), *Spodoptera frugiperda* (J. E. Smith), and banded cucumber beetle, *Diabrotica balteata* LeConte, are major maize pests causing severe economic losses [3–10]. Global annual losses to invasive pests, including these species, are estimated at US$77 billion, with FAW alone causing 40-45% yield losses in maize [8,11,12], and threatening the food security of millions of smallholder farmers worldwide [7,13–17]. The management of *S. frugiperda* and *D. balteata* has traditionally relied on synthetic pesticides, which have become ineffective due to pest resistance to many pesticide classes [7,10,18–23]. Moreover, most pesticides are classified as highly hazardous, and their application poses risks to the environment and human health [24]. With the growing consumer demand for pesticide-free food and stricter international food safety regulations [25–30], it has been recognised that a sustainable pest control strategy is required [31].

One promising alternative to synthetic pesticides is the use of entomopathogenic nematodes (EPNs) and their symbiotic bacteria. EPNs, commonly used against root herbivores, are now used against foliar pests [32–34]. EPNs from the genera *Steinernema* and *Heterorhabditis* are obligate lethal pathogens of insects. They establish mutualistic relationships with *Xenorhabdus* and *Photorhabdus* bacteria, respectively. After penetrating a suitable host, nematodes release their symbionts, which produce toxins and digestive enzymes that kill and pre-digest the insect, as well as antibiotics and repellent compounds that deter competitors, predators, and scavengers [35–44]. The nematodes and bacteria proliferate within the cadaver until resources are exhausted, then re-associate and emerge in search of new hosts, leaving behind insect remains with bacterial cells and metabolites in the soil.

In agricultural settings, EPNs and their symbionts are applied in two main forms: i) direct application of the EPNs in the soil, rhizosphere, or in aerial parts of the plant, and ii) *application of isolated* bacteria *without the nematode vector by foliar spraying or soil drenching, thanks to their high oral and contact toxicity for insects* [33,45–54]. This introduces massive amounts of bacterial cells and metabolites into the soil. *Photorhabdus* bacteria produce several toxins and metabolites with antibacterial, insecticidal, cytotoxic, antimicrobial, antifungal, acaricidal and antiparasitic properties [38,55–65]. However, the effects of *Photorhabdus* cells and metabolites on plants, soil ecosystem and non-target organisms are still unknown [66–68]. We investigated the impact of conditioning soil with i) mechanically killed or *Photorhabdus*-infected *S. frugiperda* insect larvae, ii) aqueous extracts of *Photorhabdus*-infected *S. frugiperda* insect cadavers, and iii) cell-free *Photorhabdus* supernatants on maize plant performance, resistance to herbivores and soil microbial community. We also conducted soil sterilisation and complementation experiments to investigate the links between *Photorhabdus*-mediated changes in soil microbial communities and plant performance. Our study uncovers multifaceted benefits of *Photorhabdus* bacteria as biological control agents that extend beyond agricultural pest management.

## Materials and methods

### Insect and insect rearing

*Spodoptera frugiperda* larvae were fed on an artificial diet (in g/Kg: soy milk 31.7, wheat germ 90, sugar 36.56, dried yeast 17.81, cholesterol 0.5, methylparaben 1, sorbic acid 2.3, ascorbic acid 2.46, vitamin B complex 0.184, chloramphenicol 0.269 and agar 27.5). Adults were kept in rearing cages with a continuous supply of water and a 10% sucrose solution, and eggs were collected twice weekly. Rearing was maintained at 25 ± 2°C. Fourth-to sixth-instar larvae were used for all soil conditioning experiments. *Diabrotica balteata* eggs were obtained weekly from Syngenta (Stein, Switzerland) and incubated at 25 ± 2°C until hatching. Larvae were fed on 4-day-old fresh maize roots, and second-instar larvae were used for the experiments.

### Soil material

Soil was collected from the Agroscope farm in Changins, Nyon, Switzerland (Latitude 46.398873, Longitude 6.233014). Several subsamples were collected at different locations across an area of about 4,000m^2^, then thoroughly mixed and stored in 120-litre weather-resistant plastic containers outdoors at the Institute of Biology, University of Neuchâtel.

### Bacterial strains

*Photorhabdus* strains used in this study (Table 1) comprised a genetically diverse set from five species selected for high efficacy against insect larvae in mortality tests. Bacteria were maintained in lysogeny broth (LB) medium at 28°C, re-plated monthly and renewed from glycerol stocks every six months.

**Table 1:** *Photorhabdus* bacteria strains used in the study.

| Species | Strain ID |
| --- | --- |
| *P. bodei* | ID81 |
|  | ID83 |
| *P. cinerea* | ID158 |
|  | ID163 |
| *P. khanii* subsp. *khanii* | ID316 |
|  | ID317 |
| *P. laumondii* subsp. *laumondii* | ID22 |
|  | ID25 |
|  | ID79 |
| *P. tasmaniensis* | ID138 |
|  | ID139 |
|  | ID323 |

### Insect infection assays

*Spodoptera frugiperda* larvae were infected with *Photorhabdus* strains by injecting 10 μL of bacterial suspension into the hemocoel using a Hamilton syringe. Bacterial suspensions were prepared by inoculating 25 mL LB medium with a single colony and incubating overnight at 28 °C and 180 rpm for 14–16 h, after which the optical density (OD_600_) was adjusted to OD_600_ = 1 with sterile water. Injected larvae were kept in plastic trays, and *Photorhabdus-*infected cadavers were collected every day to condition the soil as described below.

### Soil conditioning with *Photorhabdus-*infected *S. frugiperda* insect cadavers

Experimental soil was prepared by mixing field soil with sand (0-4 mm) at a 1:1 ratio. Desired amounts of experimental soil were transferred to plastic containers where mechanically killed or *Photorhabdus-*infected insect cadavers were buried equidistantly (Fig. 1). Five mechanically killed or *Photorhabdus-*infected insect cadavers per 365g of soil (equivalent to soil filled in 0.5 L plastic pots) were used for soil conditioning and kept in the greenhouse for seven days with regular moistening. Control soil was treated similarly, with no insect larvae. After seven days, conditioned soil was homogenised and transferred to 0.5 L square plastic pots to plant maize (var. Delprim). Plant growth and biomass accumulation were evaluated as described below. The experiment was repeated seven times with 10 replicates per experiment. Conditioned soil samples were collected from each experiment and stored at −20°C for soil physicochemical and microbial analyses.

**Figure 1.**
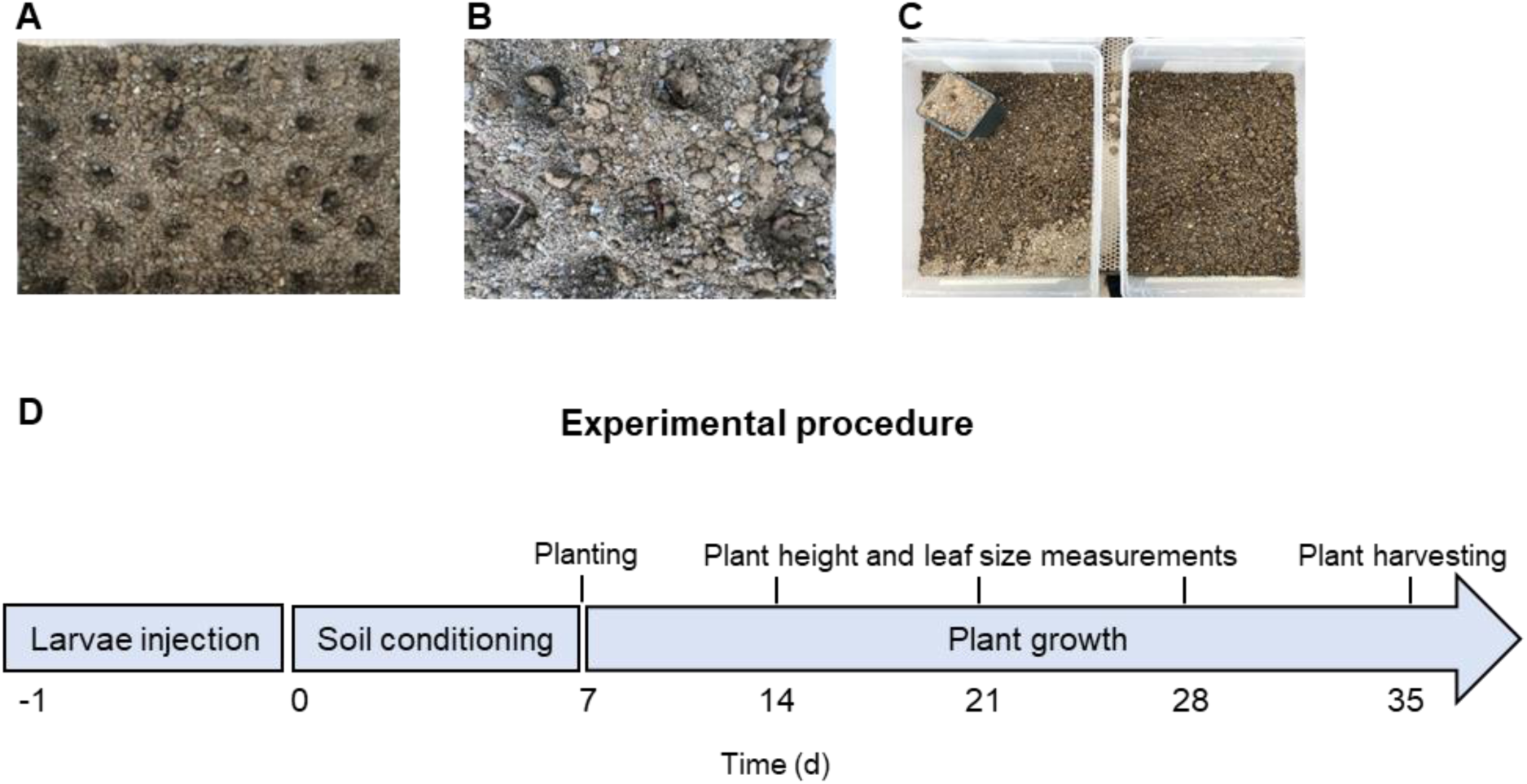
Experimental procedure used for soil-conditioning treatments. **(A)** Holes are created on the soil surface to bury *Photorhabdus*-infected or mechanically killed insect larvae. **(B)** Insect cadavers are placed in holes. **(C)** Insect cadavers were buried, and water was sprinkled on the soil surface. **(D)** Experimental time from larvae injection in the laboratory to the final plant trait measurements under greenhouse conditions.

### Soil physicochemical properties

Soil physicochemical properties were analysed at the Laboratory of Functional Ecology, University of Neuchâtel. Soil samples were oven-dried at 35 – 40 °C for 48 hours and sieved (<2 mm) to exclude stones and plant material, and subsamples of 2.5–2.55 g were used to measure bioavailable phosphorus. Sieved soil was ground into fine powder, and 10-15 mg was used for CHN analyses. We measured i) Bioavailable phosphorus (P) using sodium bicarbonate extraction (Olsen method); ii) Soil organic matter (SOM) by loss of ignition at 450 °C for 4 hours and iii) Organic carbon – Nitrogen ratio (CN) using an elemental analyser (Flash 2000, CHN-O Analyser, Thermo Scientific, Waltham, USA). Two independent soil samples per treatment were analysed.

### Soil conditioning with aqueous extracts of *Photorhabdus-*infected *S. frugiperda* insect cadavers

Eighty *Photorhabdus-*infected or mechanically killed insect larvae (≈22.5 mg each) were macerated in 18 mL of autoclaved water equivalent to the weight of larvae and filtered using 20 µm syringe filters. The resulting extracts were added to 5.8 Kg of soil and thoroughly homogenised. The conditioned soil was maintained in the greenhouse with regular moistening for seven days (Fig. 1). The control soil was treated with an equivalent volume of autoclaved water. Conditioned soils were maintained in the greenhouse for seven days with regular moistening, then re-homogenised and transferred to 0.5 L pots to grow maize (var. Delprim). The experiment was repeated three times with 10 replicates per experiment, and plant growth and biomass accumulation were evaluated as described below.

### Soil conditioning with cell-free *Photorhabdus* culture supernatants

To condition soils with cell-free *Photorhabdus* culture supernatants, 25 ml of LB medium was inoculated with a single bacterial colony and incubated at 28 °C and 180 rpm agitation for 14-16 hours. Cultures were centrifuged at 20,000 rpm for 5 minutes, and 18 mL of the cell-free supernatant was added to 5.8 kg of experimental soil and homogenised. Control soil was treated with an equivalent volume of autoclaved water. Conditioned soils were maintained in the greenhouse for seven days with regular moistening, then re-homogenised and transferred to 0.5 L pots to grow maize (var. Delprim). The experiment was repeated three times with 10 replicates per experiment, and plant growth and biomass accumulation were evaluated as described below.

### Soil sterilisation and re-inoculation experiments

To evaluate whether soil-conditioning mediated changes in soil microbial communities impact plant growth and biomass accumulation, we conducted soil sterilisation and re-inoculation experiments as follows. In the first phase, we conditioned soil with mechanically killed or *Photorhabdus*-infected insect larvae as described above. In the second phase, we autoclaved non-conditioned soil and re-inoculated soil microbial communities by mixing it with 10% of previously conditioned soil in the first phase. This procedure has been shown to re-establish natural microbial communities [69,70]. The resulting soils were left under the greenhouse for 7 days before being used for experiments. Plants were then grown on two soil types: i) non-autoclaved, non-conditioned control soil, and ii) autoclaved soil mixed with 10% of soil conditioned with mechanically killed or *Photorhabdus*-infected insect larvae. The experiments were conducted three times, with 10 replicates each time, and plant biomass accumulation was evaluated as described below.

### Plant growth measurements

To evaluate the impact of the different soil conditioning treatments on plant biomass accumulation, we harvested four-week-old plants, separated the shoots into leaves and stems and detached their roots using scissors. We gently washed the roots to remove adhering soil and placed them on tissue paper to dry. We then measured shoot and root biomass using a digital weighing scale.

### Soil DNA extraction

DNA was extracted from 200 mg of soil using TGuide S96 Magnetic Universal DNA Kit (DP812, Tiangen Biotech, Beijing, China), following the manufacturer’s recommendations. The concentration and purity of extracted DNA were assessed using a Nanodrop 2000 spectrophotometer (Thermo Fisher Scientific, Wilmington, USA). The following primers were used for amplicon sequencing: i) bacterial universal primer set 338F (5’-ACTCCTACGGGAGGCAGCA-3’) and 806R (5’-GGACTACHVGGGTWTCTAAT-3’) that amplify the V3-V4 region of 16S rRNA, ii) fungal primer set ITS1-F (5’-CTTGGTCATTTAGAGGAAGTAA-3’) and ITS2-R (5’-GCTGCGTTCTTCATCGATGC-3’) that amplify ITS1 region, and iii) nematode primer set NF1 (5′-GGTGGTGCATGGCCGTTCTTAGTT-3’) and 18Sr2b-ad (5′-TACAAAGGGCAGGGACGTAAT-3’) that amplify nematode nf1 region. Both forward and reverse primers were ligated with sample-specific Illumina index sequences. After amplification, PCR products were analysed by LabChip GX (Perkin Elmer, Waltham, USA) for fragment analysis and integrity assessment. The qualified library was sequenced on the Illumina Novaseq 6000 platform (Illumina, San Diego, USA) with paired-end 250 bp (PE250) mode. Library construction and sequencing were performed at Biomarker Technologies (BMKGENE) GmbH. To obtain Clean Reads without primer sequences, raw reads were filtered using Trimmomatic v0.33 [71] and primer sequences were removed using cutadapt 1.9.1 [72]. The DADA2 [73] method in QIIME2 2020.6 [74] was used for denoising, double-end sequence splicing, and removing chimeric sequences to obtain the final valid data (Non-chimeric Reads). Finally, using SILVA v138 as a reference database for bacteria and nematodes, and FASTQ release from UNITE v9.0 for fungi, the naive Bayes classifier combined with the alignment method was used to perform taxonomic annotation on feature sequences. Species classification information for each feature was obtained, and the community composition of each sample was statistically analysed at each level (phylum, class, order, family, genus, species). At each taxonomic level, species abundance tables were generated using QIIME software [74], and community structure diagrams were drawn in the *vegan* R package [75].

### Herbivore performance

To evaluate the impact of soil conditioning on plant resistance, *S. frugiperda* and *D. balteata* larvae were fed on 10-day-old maize plants grown on the soil conditioning treatments explained above. Ten second-instar *D. balteata* larvae were placed in solo cups and presented with fresh roots from plants grown on conditioned soil, then covered with a thin layer of soil. Five independent replicates were set up per soil conditioning treatment. Fresh roots were added every two days for ten days, and larvae were counted and weighed on the 8^th^ and 10^th^ days. For *S. frugiperda*, second-instar larvae were placed in Petri dishes and presented with leaves from plants grown on conditioned soil. Fresh leaves were added every day for four days, and larvae were counted and weighed every day. Experiments were conducted three times, with five replicates and ten larvae per replicate.

### Plant metabolomics

To evaluate plant metabolic responses to soil conditioning, we performed untargeted metabolomics on 10-day-old maize plants grown on non-conditioned control soils and soils conditioned with mechanically killed or *Photorhabdus*-infected insect larvae (n = 5 per treatment). To collect metabolomics samples, plants were removed from pots, and roots were washed to remove adhering soil. The roots and leaves were separated, wrapped in aluminium foil, flash frozen in liquid nitrogen, and stored at −80 °C for 72 hours. Flash frozen samples were ground to a fine powder under liquid nitrogen, and 50 mg of powder was weighed in a 2 mL safe-lock Eppendorf tube, followed by adding 4-8 glass beads (diameter 2-3 mm) and 1 mL of methanol:water:formic acid buffer (25:75:0.1 v/v/v) to the powder. Samples were extracted in a Qiagen tissue lyser for 3 min at 30 Hz, followed by centrifugation at 14,000 × g for 3 min. Supernatants (400 μL) were recovered in a new Eppendorf tube and stored at −80°C. Extracts were transferred to an HPLC vial and analysed by ultra-high-performance liquid chromatography-quadrupole time-of-flight tandem mass spectrometry (UHPLC-QTOFMS/MS) [76]. The system was composed of an Acquity UPLC I-Class (Waters) coupled to a Synapt XS QTOF (Waters) and controlled by MassLynx version 4.2. Chromatographic separation was performed on an Acquity UPLC™ HSS T3 column (100 × 2.1 mm, 1.8 μm, Waters) maintained at 25 °C, operating at a 0.5 mL/min flow rate. The mobile phases comprised water with 0.05% formic acid (solvent A) and acetonitrile with 0.05% formic acid (solvent B). A segmented gradient was applied, increasing from 0% to 45% B over 6.5 minutes, followed by a transition to 100% B over 3.5 minutes. The column was washed at 100% B for 3 min at 0.7 mL/min flow rate, then re-equilibrated at 0% B for another 3 min at 0.5 mL/min, with an injection volume of 1 μL. Mass spectrometry (MS) detection was performed using negative electrospray ionisation and data-dependent acquisition (DDA) mode. We selected negative ionisation as it generated richer chromatograms than positive ionisation. The Synapt XS was set at a resolution of 20,000 (at *m/z* 554), and data were collected in profile mode. The capillary and cone voltages were set at −1.0 kV and −25 V, respectively. The source and desolvation temperatures were set at 140 °C and 500 °C, respectively. The gas flow rates for desolvation, cone and collision gas (argon) were set at 1000 L/h, 250 L/h, and 2.0 L/min, respectively. The settings for DDA included a mass range for MS1 from 50 to 1200 Da, with a scan time of 0.1 seconds. The top seven MS/MS were selected, each with a 0.05-second scan time. The intensity threshold was set to 15,000 counts per second, and the MS/MS selection window was 4 Da. Peak deisotoping was activated, with dynamic exclusion set to 2.0 seconds after acquisition. An exclusion list containing the 150 most intense background ions was generated from a blank sample run prior to the sample analysis. A ramped collision energy was applied in MS/MS acquisition, ranging from 5–40 V at m/z 50 to 20–70 V at m/z 1200. Quality control samples were prepared by pooling aliquots from all samples, which were run 10 times before the sample batch and approximately every 16 samples during the batch.

Raw profile data were centroided and further converted to MzML using the microapp DataConnect (Waters). We used MZmine 4.5 for feature detection, deconvolution, and alignment of DDA (data-dependent acquisition) data, with noise thresholds set to 3000 for MS1 and 60 for MS2 [77]. The resulting peak table and MS/MS spectral data were exported and annotated using SIRIUS 6.2 [78], including CSI:FingerID for structural identification and CANOPUS for the prediction of compound classes, superclasses, and biosynthetic pathways. Taxonomically informed reannotation of molecular features was performed using the Tima R package [79] and chemical diversity clusters were visualized in Cytoscape.

### Statistical analyses

Soil chemical profiles, plant growth and insect performance were statistically analysed by analysis of variance (ANOVA) on SigmaPlot 16.0. Pairwise multiple comparisons were performed using the Holm-Sidak method (*p* ≤ 0.05). A one-sample *t*-test was conducted to determine whether the percentage change in plant biomass or larval weights (relative to controls) differed from zero (mean hypothesis; μ = 0). Soil microbial communities were analysed in R using the *vegan* package [75]. Differences in soil microbial community composition between samples were evaluated by Principal Coordinate Analysis (PCoA) and Permutational multivariate analysis of variance (PERMANOVA) on Bray-Curtis distances.

UHPLC-QTOFMS data were initially explored by a Principal Component Analysis (PCA). A Partial Least-Squares Discriminant Analysis (PLS-DA) was then performed to compare plant metabolic responses under different soil conditioning treatments and identify discriminant metabolites in resistant and susceptible plants. Cross-validation of PLS-DA models was performed using quality assessment (*Q*^2^) and R-squared (*R*^2^) parameters. The features that were upregulated in resistant and absent in susceptible and control plants were selected for metabolite identification. PCA and PLS-DA were produced using the mixOmics packages in R 4.4.2 [80].

## Results

### Conditioning soils with *Photorhabdus*-infected *S. frugiperda* insect cadavers has little effect on soil chemical properties

To test whether the different soil conditioning treatments impact soil chemical properties, we measured bioavailable phosphorus (P), organic nitrogen (N), carbon (C), and soil organic matter (SOM) in non-conditioned control soils and in soil conditioned with mechanically killed or *Photorhabdus*-infected insect larvae. The means of P, N, C, and SOM in conditioned soil varied in a treatment-specific manner but were not significantly different from those in control soils (Fig. S1). Hence, *Photorhabdus*-specific factors have little effect on soil chemical properties.

### Conditioning soils with *Photorhabdus*-infected *S. frugiperda* insect cadavers often improves plant growth

To evaluate whether *Photorhabdus*-infected insect cadavers impact plant growth through soil legacy effects, we conducted greenhouse experiments and compared biomass accumulation in maize plants grown in non-conditioned soils and in soils conditioned with either mechanically killed insect larvae or *Photorhabdus*-infected cadavers (Fig. 1). In seven independent experiments, plant biomass production was impacted by the different soil conditioning treatments in an experiment-and treatment-specific manner and independently of whether the insects used to condition the soils were infected by *Photorhabdus* or not (Fig. 2A and Fig. S2). Neutral or positive effects on plant biomass were more often observed than negative effects (Fig. 2A and Fig. S2). When all the experiments were evaluated together, plants grown on conditioned soils produced between 10 and 20% more plant biomass relative to controls (Fig. 2A). In only very few cases, we observed a significant reduction in plant biomass, which occurred in experiments 4 and 7 in a treatment-specific manner (Fig. S2). Hence, insect biomass, independently of their previous bacterial infection status, influences plant biomass accumulation through soil legacy effects.

**Figure 2.**
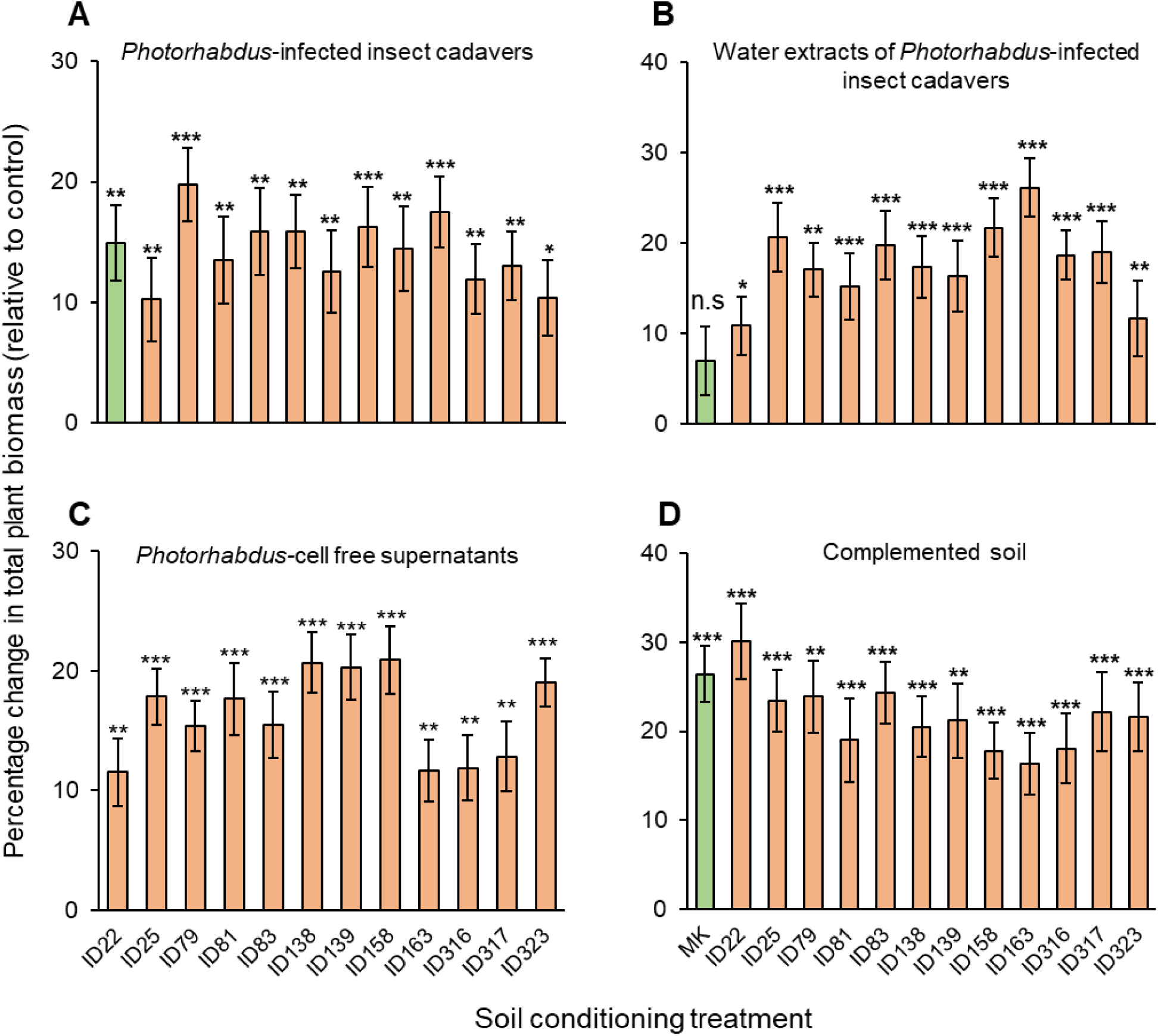
Plants accumulate more biomass when grown on conditioned soils. Percentage change (relative to controls) (± SE) in total biomass of plants grown in soil conditioned with: **(A)** *Photorhabdus*-infected insect cadavers. **(B)** Aqueous extracts of *Photorhabdus*-infected insect cadavers. **(C)** *Photorhabdus* cell-free supernatants. **(D)** autoclaved soil mixed with soil previously conditioned with *Photorhabdus*-infected insect cadavers. MK: mechanically killed larvae. IDs 22–323 refer to the different *Photorhabdus* strains used for larval infections (Table 1). The experiment in panel **A** was conducted seven times with 10 replicates each time. Experiments in panels **B-D** were conducted three independent times with 10 replicates each time. Asterisks indicate significant increases in plant biomass (*: *p* < 0.05, **: *p* < 0.01, ***: *p* < 0.001; one-sample *t*-test). n.s. not statistically significant.

### Conditioning soils with aqueous extracts of *Photorhabdus*-infected *S. frugiperda* insect cadavers improves plant growth

To test whether conditioning soil with aqueous extracts of *Photorhabdus*-infected insect cadavers influences plant growth, we compared biomass accumulation of maize plants grown in non-conditioned control soils and in soils conditioned with aqueous extracts of mechanically killed or *Photorhabdus*-infected insect larvae. In three independent experiments, neutral or positive effects of soil conditioning on plant biomass were observed (Fig. 2B and Fig. S3). When all experiments were evaluated together, plant biomass in soil conditioned with aqueous extracts of *Photorhabdus*-infected insect cadavers was increased by 11 – 26% relative to controls (Fig. 2B). In soil conditioned with aqueous extracts of mechanically killed larvae, plants gained 6.9% more biomass, which was statistically not significant relative to controls. Hence, in addition to insect biomass, *Photorhabdus*-specific factors influence plant biomass accumulation through soil-legacy effects.

### Conditioning soils with *Photorhabdus* cell-free supernatants improves plant growth

To test whether conditioning soil with *Photorhabdus* cell-free supernatants influences plant growth, we compared biomass accumulation of maize plants grown in non-conditioned control soils and in soil conditioned with *Photorhabdus* cell-free supernatants. In three independent experiments, neutral or positive effects of the different soil conditioning treatments on plant biomass were observed (Fig. 2C, Fig. S4). When all independent experiments were evaluated together, plants grown on soils conditioned with *Photorhabdus* cell-free supernatants produced 12 – 21% more biomass relative to controls (Fig. 2C). Thus, *Photorhabdus*-specific factors influence plant biomass accumulation through soil-legacy effects.

### Soil microbiota re-inoculation recapitulates the plant growth-promoting effects of *Photorhabdus*-conditioned soils

To evaluate whether *Photorhabdus*-specific effects on plant growth are linked to changes in soil microbial communities, we compared biomass accumulation of maize plants grown on non-autoclaved, non-conditioned control soil and on autoclaved soil mixed with 10% of live soil previously conditioned with either mechanically killed larvae or *Photorhabdus*-infected insect cadavers. In three independent experiments, plant biomass production was treatment-specific, with neutral or positive effects observed more frequently than negative effects (Fig. 2D and Fig. S5). Reduced plant biomass was observed only twice, in experiment 3 in autoclaved soil mixed with 10% soil conditioned with insect killed by *Photorhabdus* strains ID158 and ID323 (Fig. S5). When all experiments were evaluated together, plants grown on conditioned soils produced 16–30% more biomass than controls, independently of their previous bacterial infection status (Fig. 2D). Hence, soil conditioning-mediated changes in soil microbial communities contribute to the observed plant growth promotion effects.

### Photorhabdus-specific factors restructure soil microbial communities

#### Microbial community composition

Soil conditioning with *Photorhabdus*-infected or aqueous extracts of *Photorhabdus*-infected insect cadavers significantly altered bacterial and nematode communities more strongly than the fungal communities. Bacterial species *Myroides odoratimimus, Serratia marcescens, Providencia rettgeri,* and *Alcaligenes faecalis* were exclusively detected in conditioned soils. In contrast, *Gaiellales sp., Gemmatimonas sp., Sphingomonas* sp., *Stenotrophobacter terrae*, and *Vicinamibacterales bacteria* were present in conditioned and non-conditioned control soils but at a significantly lower abundance in soil conditioned with *Photorhabdus*-infected or aqueous extracts of *Photorhabdus*-infected insect cadavers *compared to controls* (Figs. 3, 4, S6, S7). Moreover, *Myroides odoratimimus*, *Serratia marcescens*, *Providencia rettgeri*, and *Alcaligenes faecalis* were 5–29% more abundant in soils conditioned with *Photorhabdus*-infected cadavers than in soils conditioned with mechanically killed larvae (1–3%) (Fig. 3). Consistently, these species were abundant in soils conditioned with aqueous extracts of *Photorhabdus*-infected cadavers but were absent in soils conditioned with aqueous extracts of mechanically killed larvae (Fig. 4). Together, these results indicate that *Photorhabdus*-specific factors significantly influenced the bacterial community.

**Figure 3.**
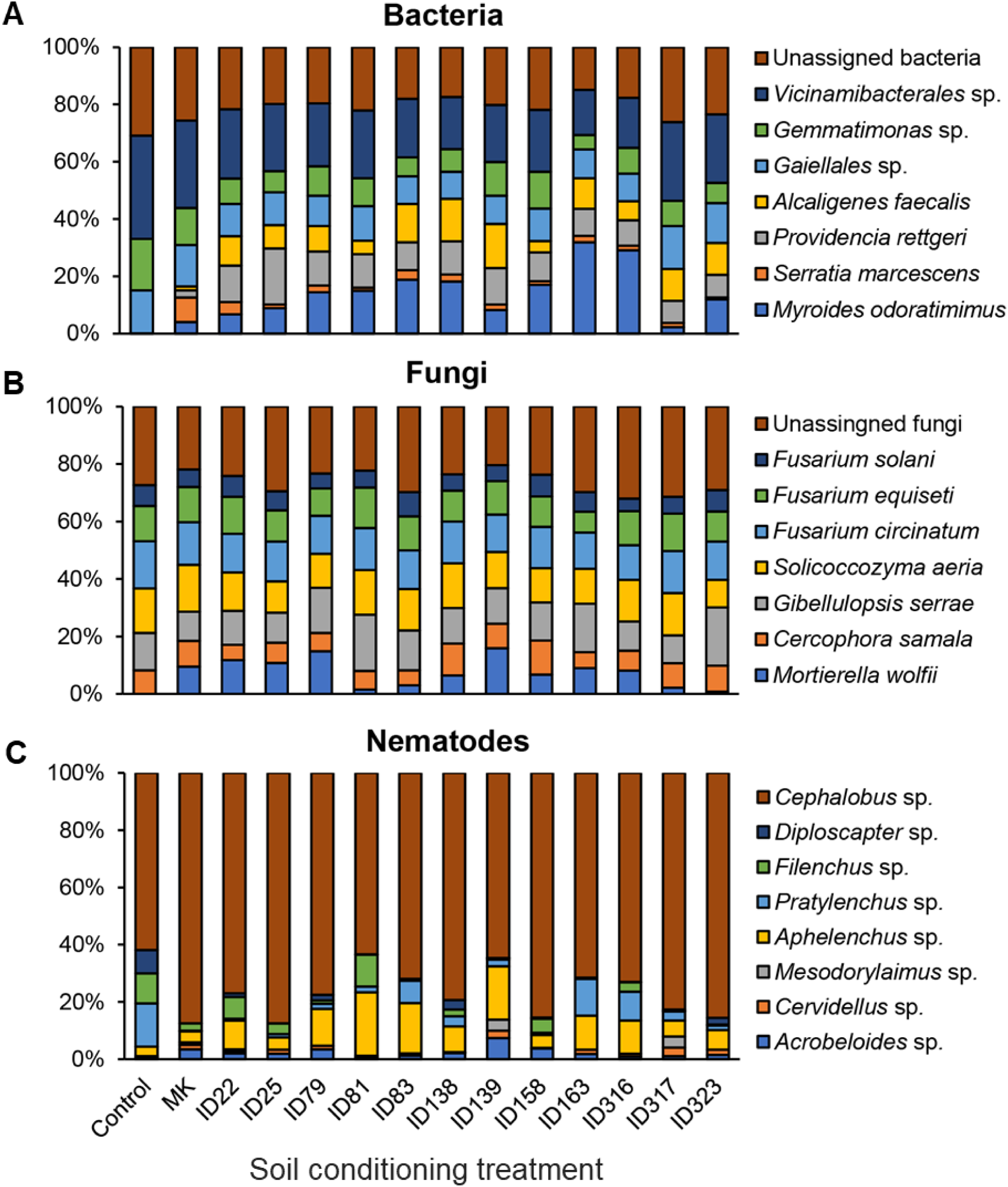
*Photorhabdus*-specific factors restructure soil microbial communities. Relative abundance of soil: **(A)** bacteria, **(B)** fungi, and **(C)** nematodes in non-conditioned control soils and in soils conditioned with mechanically killed larvae (MK) or *Photorhabdus*-infected insect cadavers. IDs 22–323 refer to the different *Photorhabdus* strains used for larval infections (Table 1).

**Figure 4.**
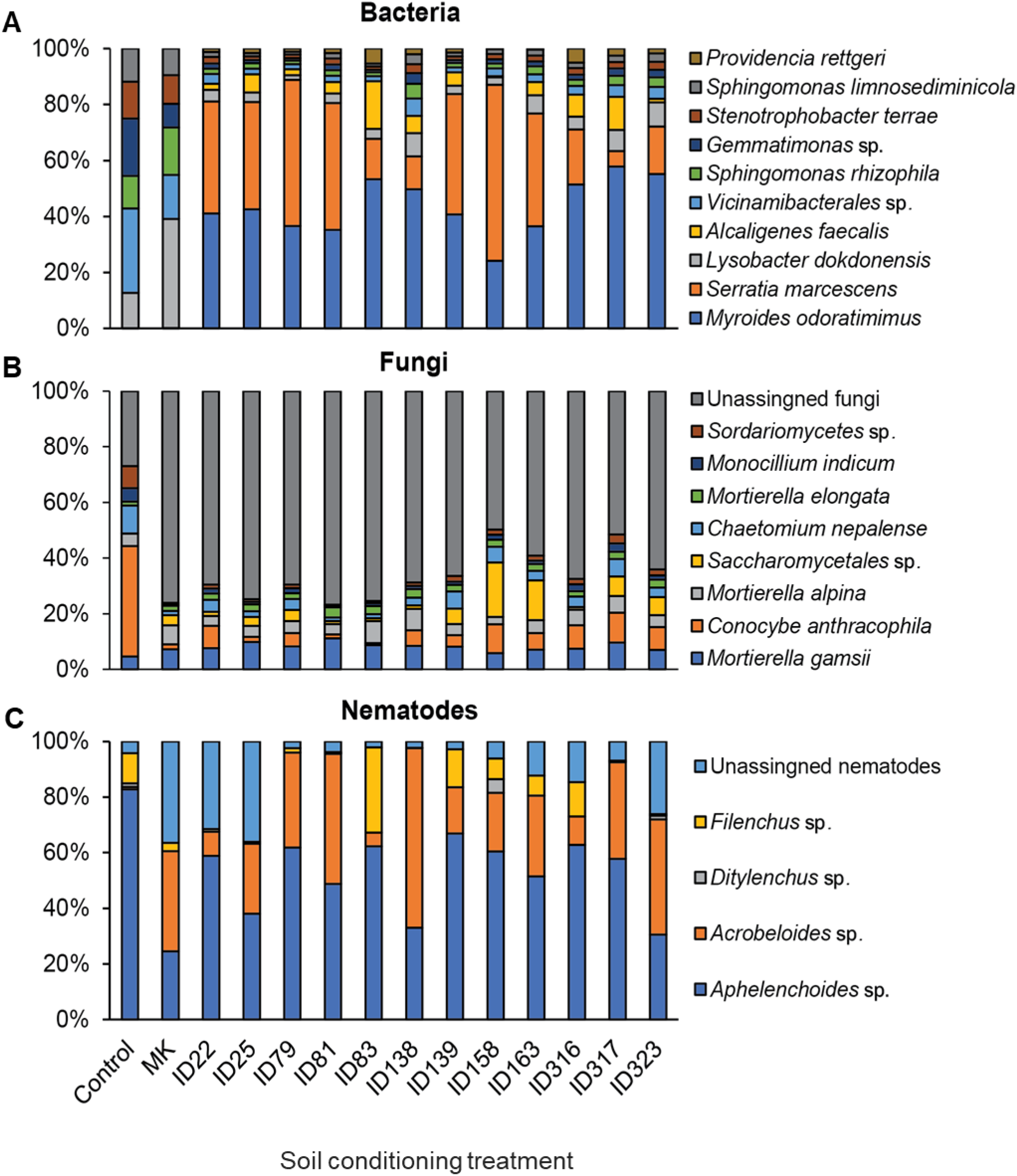
*Photorhabdus*-specific factors restructure soil microbial communities. Relative abundance of soil: **(A)** bacteria, **(B)** fungi, and **(C)** nematodes in non-conditioned control soils and in soils conditioned with water extract of mechanically killed larvae (MK) or *Photorhabdus*-infected insect cadavers. IDs 22–323 refer to the different *Photorhabdus* strains used for larval infections (Table 1).

The fungal community in soils conditioned with *Photorhabdus*-infected or mechanically killed larvae did not show significant differences relative to non-conditioned control soils. All detected fungal species, including *Mortierella wolfii, Cercophora samala, Gibellulopsis serrae, Solicoccozyma aeria, Fusarium circinatum, Fusarium equiseti,* and *Fusarium solani* were present in both conditioned and non-conditioned control soils, with no significant differences in their abundance (Figs. 3 and S8). Contrary to our expectations, the fungal community was significantly altered in soils conditioned with aqueous extracts of *Photorhabdus*-infected insect cadavers. The fungal species, *Mortierella gamsii, Mortierella alpina, Mortierella elongata, Saccharomycetales,* and unassigned fungi were significantly more abundant, while *Conocybe anthracophila, Chaetomium nepalense, Monocillium indicum,* and *Sordariomycetes* sp. were significantly less abundant in soils conditioned with aqueous extracts of *Photorhabdus*-infected *S. frugiperda* insect cadavers compared to non-conditioned control soils (Fig. 4 and S9).

In the nematode community, *Cephalous sp., Aphelenchus sp., Pratylenchus sp., Filenchus sp., Acrobeloides sp., Cervidellus sp., Diploscapter sp.,* and *Mesodorylaimus sp. were* detected in soils conditioned with *Photorhabdus*-infected insect cadavers (Fig. 3). Notably, *Cephalous sp., Aphelenchus sp., Acrobeloides sp., Cervidellus sp.,* and *Mesodorylaimus sp.* were more abundant and exclusively detected in conditioned soils but not in control soil. Only *Cephalous sp. was detected in both* conditioned *and non-conditioned control soil.* Additionally, the abundance of *Cervidellus sp., Mesodorylaimus sp., filenchus sp., Diploscapter sp. and Pratylenchus sp. in* conditioned soil was treatment-specific (Fig. S10). In soils conditioned with aqueous extracts of *Photorhabdus*-infected insect cadavers, *Aphelenchoides* sp., *Acrobeloides* sp., *Ditylenchus* sp., *Filenchus* sp., and unassigned nematodes were detected (Fig. 4). However, only *Acrobeloides* sp and unassigned nematodes were more abundant in conditioned soil than in non-conditioned control soil (Fig. S11).

#### Microbial diversity

To investigate the effect of soil conditioning on microbial diversity, we compared microbial communities across treatments using PCoA and PERMANOVA based on Bray-Curtis distances. We observed that soil conditioning with *Photorhabdus*-infected or with aqueous extracts of *Photorhabdus*-infected insect cadavers significantly altered the bacterial communities. The PCoA analysis showed a higher dissimilarity index in bacterial communities across soil conditioning treatments compared to non-conditioned control soil. Additionally, the PERMANOVA test confirmed a significant effect of soil conditioning on the bacterial community composition (Fig. S12 and S13). Similar significant changes were observed with the fungal and nematode communities in soils conditioned with aqueous extracts of *Photorhabdus*-infected insect cadavers (Fig. S13). In soil conditioned with *Photorhabdus*-infected insect cadavers, PCoA analyses showed no dissimilarity in the fungal community, and the PERMANOVA revealed no significant effects of soil conditioning on the fungal community composition (Fig. S12). Similarly, no significant differences in the nematode community were observed (Fig. S12). However, individual nematode species abundance was significantly altered by soil conditioning relative to controls (Fig. S10). Overall, we demonstrate that conditioning soils with mechanically killed or *Photorhabdus*-infected insect larvae alters soil bacterial and nematode communities, and to a lesser extent, fungal communities.

### Conditioning soils with *Photorhabdus*-infected insect cadavers often trigger plant resistance to herbivores in a *Photorhabdus* strain-specific manner

To test whether conditioning soils with *Photorhabdus*-infected insect cadavers influences plant resistance against insect herbivores, we compared the performance of *D. balteata* and *S. frugiperda* larvae fed on maize plants grown on non-conditioned control soils and on soils conditioned with mechanically killed or *Photorhabdus*-infected insect larvae. Across three independent experiments, neutral, positive and negative effects on insect performance were observed in a *Photorhabdus* strain-specific manner (Fig. 5A and S14). When the three experiments were evaluated together, plants growing in soils conditioned with cadavers killed by strains ID139, ID158, ID163, ID79, and ID323 were more resistant to *D. balteata* larvae, reducing their weights by 15 – 20% relative to controls (Fig. 5A). Neutral effects were observed in larvae that fed on plants grown on soils conditioned with mechanically killed and insects killed by *Photorhabdus* strains ID22, ID25, ID81, ID83, ID138, ID317, and ID316. Thus, the larval weights were not statistically significant (Fig. 5A).

**Figure 5.**
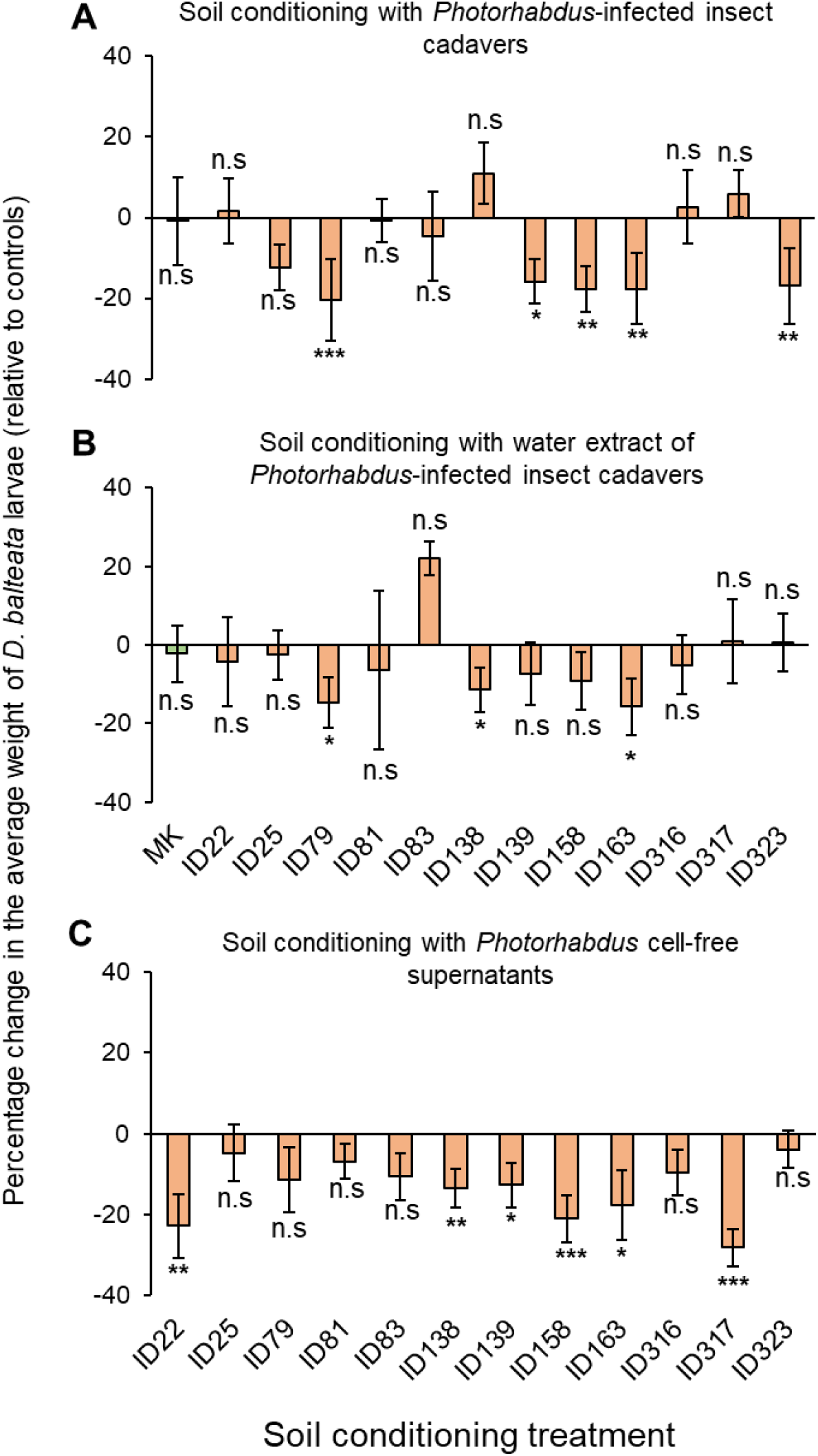
Plants grown on conditioned soils often resist the attack of *D. balteata*. Percentage change (relative to controls) (± SE) in average weight of *D. balteata* larvae fed with plants grown in soil conditioned with: **(A)** *Photorhabdus*-infected insect cadavers. **(B)** aqueous extracts of *Photorhabdus*-infected insect cadavers. **(C)** *Photorhabdus* cell-free supernatants. MK: mechanically killed larvae. IDs 22–323 refer to the different *Photorhabdus* strains used for larval infections (Table 1). These experiments were conducted three independent times, with 5 replicates each time and 10 larvae per replicate. Asterisks above bars indicate significant changes in the average weight of *D. balteata* larvae (*: *p* < 0.05, **: *p* < 0.01, ***: *p* < 0.001; one-sample *t*-test). Positive values indicate weight gain and negative values indicate weight lost. n.s. not statistically significant.

For *S. frugiperda* larvae feeding on maize leaves, we observed neutral effects, except for plants grown on soils conditioned with insects killed by *Photorhabdus* strains ID158, ID22, and ID138, which reduced larval weights by 10 – 59% relative to controls (Fig. 6A and S15). Notably, only plants grown on soils conditioned with insects killed by *Photorhabdus* strain ID158 significantly suppressed the feeding of both *D. balteata* and *S. frugiperda* larvae (Fig. 5A and 6A). The feeding of both *herbivores* on plants grown on soil conditioned with mechanically killed insect larvae was not affected, with no significant differences in larval performance (Figs. 5A, 6A, S14 and S15).

**Figure 6.**
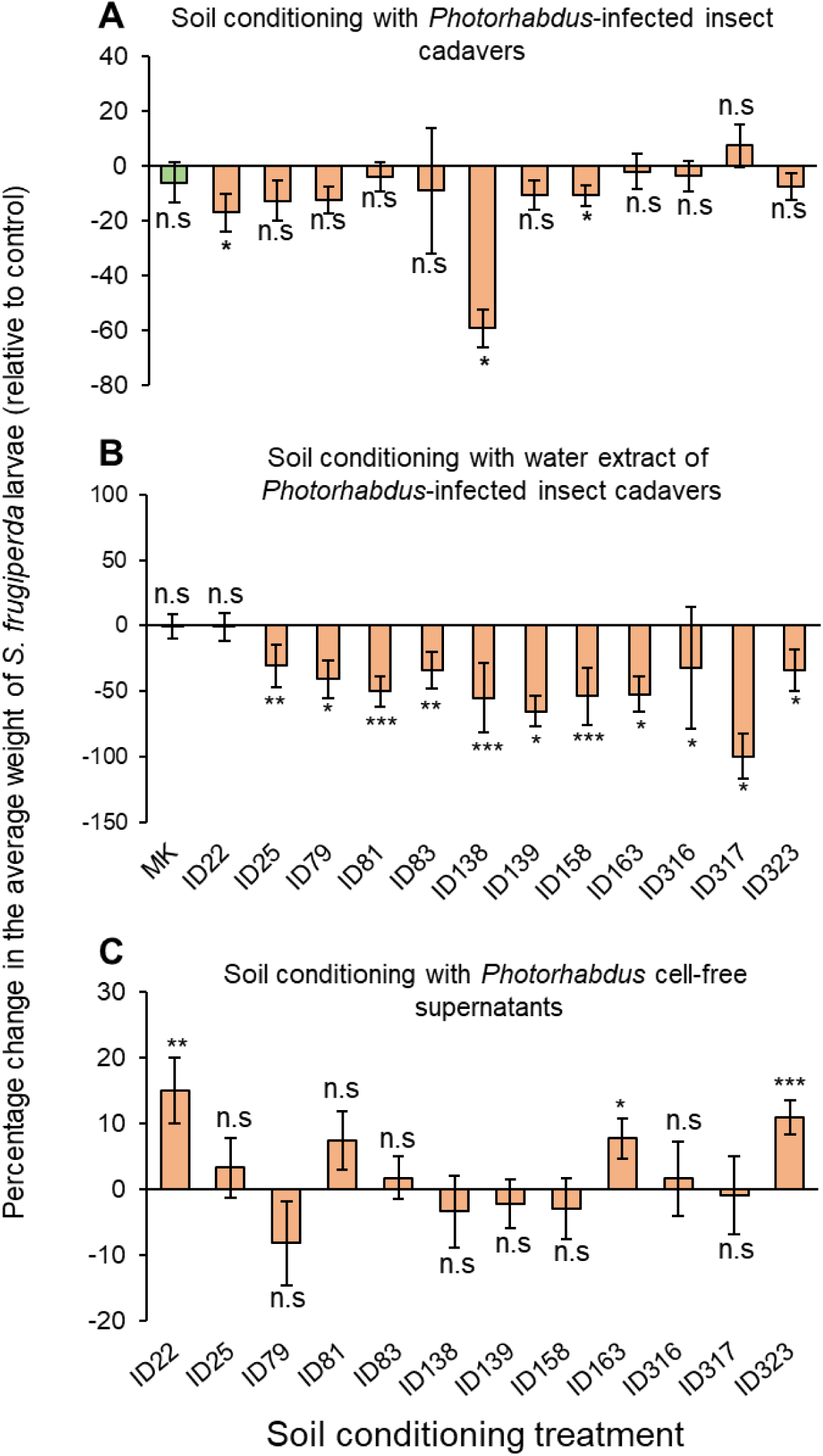
Plants grown on conditioned soils often resist the attack of *S. frugiperda*. Percentage change (relative to controls) (± SE) in average weight of *S. frugiperda* larvae induced by plants grown in soil conditioned with: **(A)** *Photorhabdus*-infected insect cadavers. **(B)** aqueous extracts of *Photorhabdus*-infected insect cadavers. **(C)** *Photorhabdus* cell-free supernatants. MK: mechanically killed larvae. IDs 22–323 refer to the different *Photorhabdus* strains used for larval infections (Table 1). These experiments were conducted three independent times, with 5 replicates each time and 10 larvae per replicate. Asterisks above bars indicate significant reductions or increases in the average weight of *S. frugiperda* larvae (*: *p* < 0.05, **: *p* < 0.01, ***: *p* < 0.001; one-sample *t*-test). n.s. not statistically significant.

### Conditioning soil with aqueous extracts of *Photorhabdus*-infected *S. frugiperda* insect cadavers often trigger plant resistance to herbivores in a strain-specific manner

Across three independent experiments, *D. balteata* larvae showed neutral, positive and negative responses when fed on plants grown in soils conditioned with aqueous extracts of *Photorhabdus*-infected insect cadavers (Fig. 5B and S16). Plants grown in soils conditioned with aqueous extracts of insects killed by strains ID79, ID138, and ID163 were significantly more resistant, reducing larval weights by 11 – 15% relative to controls (Fig. 5B). Neutral effects were observed in larvae that fed on plants grown on soils conditioned with aqueous extracts of mechanically killed or insects killed by the remaining *Photorhabdus* strains, with no significant differences observed (Fig. 5B and S16).

For *S. frugiperda*, we observed negative and neutral, rather than positive effects on larval performance (Fig. 6B and S17). Plants grown on all conditioned soils resisted the attack of *S. frugiperda* larvae, significantly reducing larval weights by 30 – 100%. Only plants grown on soils conditioned with aqueous extracts of mechanically killed or insects killed by *Photorhabdus strain ID22 did* not affect larval feeding (Fig. 6B). These results show that *Photorhabdus*-specific factors can induce plant resistance against root and leaf-feeding herbivores.

### Conditioning soils with *Photorhabdus* cell-free supernatants triggers plant resistance against root, but not leaf-feeding herbivores

Across three independent experiments, we consistently observed neutral and negative effects on insect performance in plants grown on soils conditioned with *Photorhabdus* cell-free supernatants, rather than positive effects (Fig. 5C and S18). When three experiments were evaluated together, all plants grown in soils conditioned with *Photorhabdus* cell-free supernatants suppressed the growth of *D. balteata* larvae, reducing their weight by 3 – 28% relative to controls (Fig. 5C and S18). However, significant resistance against *D. balteata* larvae was observed in plants grown in soils conditioned with supernatants of *Photorhabdus* strains ID22, ID138, ID139, ID158, ID163, and ID317, which reduced larval weight by 12–28% relative to controls (Fig. 5C and S18).

For *S. frugiperda*, we observed negative and neutral, rather than positive effects, irrespective of the *Photorhabdus* strain (Fig. 6C and S19). Plants grown in soils conditioned with supernatants of *Photorhabdus* strains ID79, ID138, ID139, ID158, and ID317 tended to resist *S. frugiperda* larvae, reducing larval weight by 1 – 8%, though not statistically significant relative to controls (Fig. 6C and S19). In contrast, plants grown on the other conditioned soils increased larval weights by 2 – 15% relative to controls, with significant weight increases (8 – 15%) observed in larvae fed on plants grown on soil conditioned with supernatants of *Photorhabdus* strains ID22, ID163, and ID323 (Fig. 6C and S19). These results show that conditioning soil with *Photorhabdus* cell-free supernatants can trigger plant resistance against root-feeding than leaf-feeding herbivores.

### Conditioning soils with insects killed by Photorhabdus induces plant metabolic responses

To evaluate plant metabolic responses to soil conditioning, we analysed low-molecular-weight metabolites of plants grown on non-conditioned control soils and soils conditioned with mechanically killed or *Photorhabdus*-infected insect cadavers. Based on insect performance results (Figures 5 and 6), plants were categorised as “Resistant” and “Susceptible”. A PLS-DA was performed to identify soil conditioning treatments that induced resistance responses in roots and leaves. We found treatment-specific changes in plant metabolites. In roots, plants grown on soils conditioned with insects killed by *Photorhabdus* strains ID139, ID158, ID163, ID79, or ID 323 exhibited distinctive root metabolites (Fig. 7A). In leaves, distinctive metabolites were observed in plants grown on soil conditioned with insects killed by *Photorhabdus* strains ID158, ID22, or ID138 (Fig. 7B). These findings are consistent with insect performance results, where plants grown on these soil conditioning treatments resisted the attack of *D. balteata* and *S. frugiperda*.

**Figure 7.**
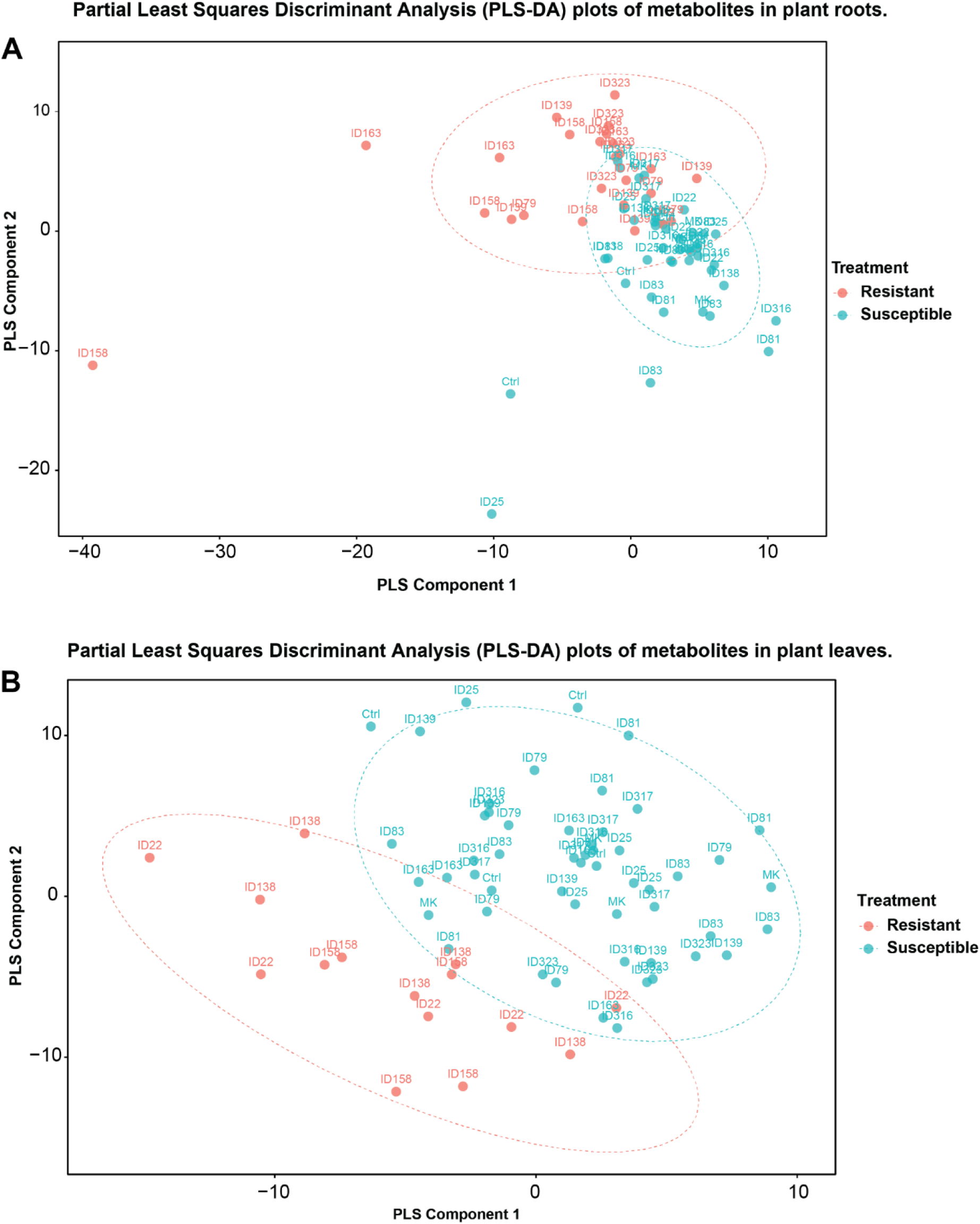
Leaves and roots of plants respond to soil conditioning treatments at the metabolic level. Partial Least-Squares Discriminant Analysis (PLS-DA) score plots of metabolites in **(A)** roots and **(B)** leaves of plants grown on non-conditioned control soils and soils conditioned with *Photorhabdus*-infected insect cadavers or with mechanically killed insect larvae (MK). IDs 22–323 denote the *Photorhabdus* strains used for larval infection (Table 1). Plants grown under the different soil conditioning treatments were classified as resistant (red) or susceptible (blue) based on insect performance results (Figures 5 and 6).

To identify metabolic changes associated with resistant plants, we focused on the most abundant metabolites highlighted in the enhanced molecular network. We observed a higher accumulation of specific metabolites in resistant plants than in susceptible plants. In roots, Twelve metabolites belonging to alkaloids, benzoxazinoids, fatty acids, shikimates, phenylpropanoids, and terpenoids were upregulated (Table S1). Four metabolites, namely Aromadendrin, Compound ID 196, Imidazole alkaloid, and (6R,9R,10R)-6,9,10-trihydroxyoctadec-7-enoic acid, were detected in both resistant and susceptible plants, but were significantly upregulated in resistant plants. The other eight metabolites, 4-[(6R)-6-hydroxy-5,5-dimethylcyclohexen-1-yl] benzoic acid, 2 cis-Abscisic Acid, Canthoside B, Apigenin 7-sulfate, Pimarane and Isopimarane diterpenoids, Ajugaciliatin A, GDIMBOA and Tinosineside B, were exclusively detected in resistant and absent in susceptible and control plants (Fig. S20). In leaves, seven metabolites belonging to amino acids, fatty acids, and phenylpropanoids were upregulated (Table S1). Four of these metabolites, (2R)-2-ammonio-3-phenylpropanoate, L-Tryptophan, 1-Acyl-sn-glycero-3-phosphoglycerol (N-C16:1), and 2 cis-Abscisic Acid, were detected in both susceptible and resistant plants, but were significantly upregulated in resistant plants (Fig. S21). The other metabolites, Cherimola cyclopeptide, PC (16:0/18:2(9Z,12Z)) 1-hexadecanoyl-2-(9Z,12Z-octadecadienoyl)-sn-glycero-3-phosphocholine, and PA (22:4(7Z,10Z,13Z,16Z) /22:2(13Z,16Z)) were exclusive to resistant plants (Fig. S21). Notably, only 2-cis-Abscisic Acid was consistently upregulated in roots and leaves (Fig. S20 and S21). Altogether, our results demonstrate that conditioning soils with *Photorhabdus*-infected insect cadavers induces metabolic responses in plant roots and leaves, with patterns that differ between resistant and susceptible plants.

## Discussions

We demonstrated that conditioning soils with mechanically killed or *Photorhabdus*-infected insect larvae, aqueous extracts of *Photorhabdus*-infected insect larvae, or cell-free *Photorhabdus* supernatants improve maize plant growth in a greenhouse environment. Our findings add more evidence to previous laboratory experiments [81], showing that *Photorhabdus* strains *P. temperata* M1021 and *P. luminescens* TT01 improve plant growth when inoculated in rice plant seedlings. This is encouraging for growers, as *Photorhabdus* bacteria could offer dual benefits of pest control and plant growth promotion in fields where they are applied [45–49,52,53]. We also observed increased plant biomass in soils conditioned with mechanically killed insect larvae, consistent with studies that used insect cadavers, frass and exuviae for soil amendment [82–84]. Insect-based materials contain a large reserve of plant-available nitrogen, which can enhance plant growth when released into the soil [85–89]. Insects weigh approximately 10% organic nitrogen and contain carbohydrates, lipids, protein and chitin [86,90], making them richer in ammonium and nitrate levels, which can have more lasting effects in the soil [85,88,91–93]. Additionally, decomposing insect materials stimulate the growth of beneficial soil rhizobacteria and accelerate their activities to break down nutrients in a plant-available form and directly enhance plant growth [86,94,95].

The soil conditioning treatment with mechanically killed insect larvae differs from soil conditioning with *Photorhabdus*-infected or aqueous extracts of *Photorhabdus*-infected insect cadavers, or cell-free supernatants in mechanisms of plant growth enhancement. Whilst mechanically killed insects directly introduce nutrients into the soil, *Photorhabdus-*infected insect cadavers are likely depleted in nutrients but full of bacterial biomass and their by-products, such as toxins and metabolites. Therefore, plant growth stimulation in soils conditioned with *Photorhabdus*-infected insect cadavers or *Photorhabdus* metabolites could be explained in two ways. One possible mechanism is the incorporation of *Photorhabdus-*produced bioactive metabolites, such as resorcinol, siderophores, indole-3-acetic acid and gibberellins into the soil [68,81,96]. These bioactive metabolites are known to stimulate plant growth upon direct interaction with plants via hormone-driven processes, resulting in accelerated growth, increased biomass and higher crop yields [97–101]. This argument was supported by the fact that conditioning soils with aqueous extract of *Photorhabdus*-infected insect cadavers or *Photorhabdus* cell-free supernatants significantly increased plant biomass, whereas soil conditioning with aqueous extracts of mechanically killed larvae had no significant effect relative to controls. These findings indicate that *Photorhabdus*-specific effects, together with insect biomass, drive the observed plant growth promotion effects. However, whilst studies have recognised microbial metabolites as plant growth promoters [102–109], we do not know the metabolic and biochemical mechanisms underlying *Photorhabdus*-mediated plant growth promotion. This would require the isolation and identification of *Photorhabdus* metabolites associated with plant growth promotion. Another mechanism behind the increased plant growth could be changes in microbial communities through *Photorhabdus*-mediated effects. To explore this mechanism, we conducted sterilisation and re-inoculation experiments to investigate the link between *Photorhabdus*-mediated changes in soil microbial communities and their soil feedback on plant growth. We demonstrated that autoclaved soils complemented with only 10% of living soil that had been conditioned with mechanically killed or *Photorhabdus*-infected insect larvae significantly increased plant total biomass relative to controls. This indicates that *Photorhabdus*-mediated changes in soil microbial communities indeed positively impact plant growth. These results agree with studies reporting that changes in soil microbial communities drive plant growth and resistance in soils inoculated with microbial agents [110–113].

We also demonstrated that conditioning soils with *Photorhabdus*-infected insect cadavers or their aqueous extracts significantly altered the bacterial, nematode and to a lesser extent, fungal communities. The findings are consistent with studies reporting changes in bacterial and nematode communities in soils inoculated with beneficial microbes [110,114,115]. For instance, inoculating soil with *Pseudomonas* species, *P. chlororaphis,* and *P. putida* restructured the bacterial community from a Gram-positive to Gram-negative dominant community in maize rhizosphere [110], while *Bacillus thuringensis* (Bt) toxins altered soil bacterial and nematode community composition [114,116–119]. Changes in soil microbial communities can positively or negatively affect plant growth [120,121]. In our study, bacterial species, *Myroides odoratimimus, Serratia marcescens, Providencia rettgeri,* and *Alcaligenes faecalis* were exclusively abundant in conditioned soils, not in non-conditioned control soil. These bacterial species enhance plant growth through nutrient cycling [122–128], improve plant stress tolerance, and *protect plants against attackers* [122,129,130]. In the nematode community, soil conditioning shifted the abundance of plant pathogenic nematodes, *Pratylenchus sp.* and *Filenchus sp.* [114,131,132] and resulted in enrichment with beneficial nematodes, including *Cephalobus sp., Aphelenchus sp., Acrobeloides sp., Cervidellus sp., Diploscapter sp.,* and *Mesodorylaimus sp.* These beneficial nematodes are bacterivorous and fungivorous [114,132–136] and known to contribute to nutrient mineralisation and suppression of plant pathogens [137,138]. Notably, *Aphelenchus sp., Acrobeloides sp., Cervidellus sp.,* and *Mesodorylaimus sp.* were exclusively enriched in conditioned soils, but their abundance was treatment dependent. Given that the production of certain metabolites varies across *Photorhabdus* strains [68,139,140], strain-specific chemical inputs may have differentially shaped nematode community composition. Additionally, the increased abundance of bacterivorous and fungivorous nematodes in certain soil conditioning treatments may reflect the availability of suitable prey hosts, which likely promoted their population growth [136,141–145]. In the fungal community, conditioning soils with aqueous extracts of *Photorhabdus*-infected insect cadavers significantly altered their composition, increasing the abundance of *Mortierella gamsii, Mortierella elongata, Mortierella alpina*, and *Saccharomycetales* sp than in non-conditioned control soils. These fungal species have been described as plant growth promoters and exert competition against plant pathogens [146–153]. However, conditioning soils with *Photorhabdus*-infected insect cadavers did not affect the fungal community. This may be because the amount of toxins released during cadaver decomposition was insufficient to induce measurable changes. Rasool et al. [154] showed that changes in soil fungal communities depend on the density of the inoculant. In their experiments, the fungal communities changed when *M. brunneum* was applied by soil drenching, which increased its density and persistence in the soil, but not seed treatment [154]. Similarly, applying microbial inoculants via seed treatment or rhizosphere application resulted in limited, transient or no significant effects on the soil fungal community [155–157], with transient effects disappearing shortly [155]. The fungal community in soil conditioned with *Photorhabdus*-infected insect cadavers included two beneficial species, *Mortierella wolfii* and *Solicoccozyma aeria* [151,158–160], and was dominated by plant pathogens, including *Gibellulopsis serrae, Fusarium circinatum, Fusarium equiseti,* and *Fusarium solani* [161–164]. *Fusarium* sp., which was most dominant in control soil, is known to reduce crop productivity by colonising plant roots and disrupting protein synthesis and water transport in plants [165,166]. Altogether, conditioning soil with *Photorhabdus*-infected insect cadavers or their aqueous extracts increased the abundance of beneficial bacteria, fungi and nematodes, which may have mitigated plant pathogen pressure, and enhanced plant growth relative to non-conditioned controls. Thus, *Photorhabdus*-specific factors influence soil microbial communities with downstream consequences for plant performance.

In this study, we further demonstrate that the performance of *D. balteata* and *S. frugiperda* larvae feeding on maize plants grown on soil conditioned with *Photorhabdus*-infected insect cadavers, aqueous extracts of *Photorhabdus*-infected insect cadavers or *Photorhabdus* cell-free supernatants is affected, either negatively, positively or neutrally, in a *Photorhabdus* strain-specific manner. These results are consistent with studies that reported suppressive effects of plants grown on soils conditioned with microbial agents against root-feeding insects [167–170] and leaf-feeding herbivores [171–175]. Such negative effects on herbivores are often reported to be mediated by changes in plant secondary metabolites following soil conditioning [170,176,177]. However, neutral or positive effects on root- and leaf-feeding insects have also been documented, with herbivore-specific responses [177–185]. Induced plant resistance affects generalist more strongly than specialist herbivores [186–188], as some insects can detoxify or neutralise plant defensive metabolites and mitigate their bioactivity [189–192]. In our study, both generalist, *D. balteata* and specialist *S. frugiperda* larvae were negatively affected. Thus, the differences in plant resistance among soil-conditioning treatments may be explained by variations in defence-related metabolites accumulated in plants. *Photorhabdus* spp. produce diverse bioactive metabolites, including trans-cinnamic acid, siderophores, resorcinol, terpenes and nucleosides, that influence plant growth and defence pathways [68,98,193–196]. As metabolite production varies among *Photorhabdus* strains [68,139,140], strain-specific differences in plant responses to herbivory are plausible. Moreover, *Photorhabdus*-mediated changes in soil microbial communities may also influence plant chemical profiles and insect performance, like in previous studies [197–200], potentially explaining why certain soil conditioning treatments resulted in a stronger plant resistance effect.

Lastly, we confirm our hypothesis that conditioning soils with *Photorhabdus*-infected insect cadavers increased the levels of maize defence metabolites, explaining the enhanced resistance to herbivores. Conditioning soils with *Photorhabdus*-infected insect cadavers significantly upregulated certain metabolites, which varied with *Photorhabdus strains. Notably, resistant plants* accumulated distinct metabolites relative to susceptible and control plants. In roots, Aromadendrin, Compound ID 196, Imidazole alkaloids, and 6,9,10-trihydroxyoctadec-7-enoic acid were more abundant in plants grown on soils conditioned with insects killed by strains ID79, ID139, ID158, ID163, and ID323. These metabolites have many biological activities and could be involved in the defence mechanisms of plants [201–204]. Additionally, only resistant plants accumulated metabolites including 4-[(6R)-6-hydroxy-5,5-dimethylcyclohexen-1-yl] benzoic acid, 2 cis-Abscisic Acid, Canthoside B, Apigenin 7-sulfate, Pimarane and Isopimarane diterpenoids, Ajugaciliatin A, GDIMBOA and Tinosineside B, many of which have their plant defensive roles against herbivores and pathogens documented [205–209]. Although Ajugaciliatin A, Canthoside B, Apigenin 7-sulfate and Tinosineside B have not yet been linked to plant defence, their biological activities, including antimicrobial, antioxidant, cytotoxic, and anti-estrogenic effects [210–214], suggest potential roles in resistance or defence signalling. In the leaves, more abundant metabolites included (2R)-2-ammonio-3-phenylpropanoate, L-Tryptophan, 1-Acyl-sn-glycero-3-phosphoglycerol (N-C16:1), 2 cis-Abscisic Acid, Cherimola cyclopeptide, PC (16:0/18:2(9Z,12Z)) 1-hexadecanoyl-2-(9Z,12Z-octadecadienoyl)-sn-glycero-3-phosphocholine and PA(22:4(7Z,10Z,13Z,16Z)/22:2(13Z,16Z)), all reported to confer resistance against insect pests and pathogens [203,215–221]. These metabolic changes are consistent with studies of plant responses to root exposure to *Photorhabdus* bioluminescence or EPN-killed insect cadavers, although the mechanisms by which plants respond to these exposures remain unresolved [44,222–224]. Root exposure to *Photorhabdus* bioluminescence induced defensive metabolites in maize and reduced *D. balteata* larval growth [44]. Similarly, exposing potato and maize plant roots to EPN-killed insect cadavers triggered systemic defence responses, deterring the feeding of *Leptinotarsa decemlineata* and *S. frugiperda* larvae [222–224]. However, *S. frugiperda* caterpillars feeding on leaves of bioluminescence-exposed plants were unaffected [44], suggesting variations in the biological effects of induced systemic responses. These metabolic changes also impact other trophic levels, influencing nematodes, insects, and plants [44,223,224]. Taken together, this implies that conditioning soils with insects killed by selected *Photorhabdus* strains can confer systemic responses in maize that enhance resistance to below and aboveground herbivores.

## Acknowledgments

The authors thank the Institute of Biology of the University of Neuchâtel (Switzerland) and the Swiss National Science Foundation for their support. The authors are thankful to Claudia Di Cesare, Amandine Pillonel, and Jouini Ilham for technical support. This study was supported by the Swiss National Science Foundation (grant 186094 to R.A.R.M. and grants 204811 and 217975 to S.R.) and the Swiss Government Excellence Scholarship (grant 2021.0663 to J.E.).

## Author contributions

Conceptualisation, R.A.R.M.

Data curation, J.E., R.A.R.M.

Formal analysis, J.E., R.A.R.M., D.E.

Funding acquisition, R.A.R.M., T.C.J.T.

Investigation, J.E., C.C.M.A., R.A.R.M.

Methodology, R.A.R.M., J.E., C.C.M.A., and G.G.

Project administration, R.A.R.M.

Resources, R.A.R.M., W.Z., S.R., and T.C.J.T.

Supervision, R.A.R.M.

Validation, R.A.R.M.

Visualisation, J.E. and R.A.R.M.

Writing – original draft, J.E. and R.A.R.M.

Writing – review & editing, R.A.R.M., J.E.

