## Supplementary material for "*Photorhabdus* metabolites reshape soil microbial communities and promote plant growth and insect resistance"

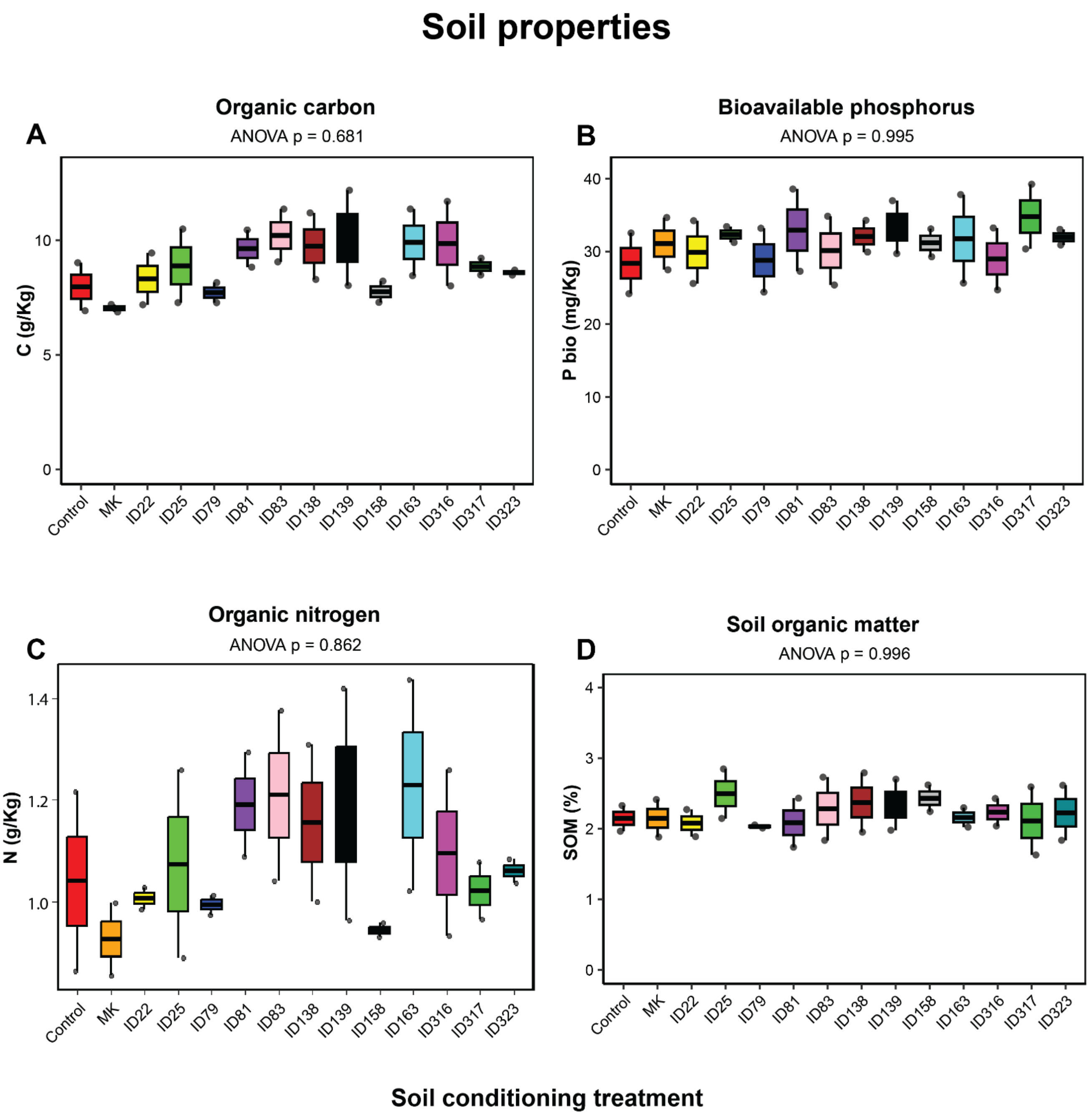


**Figure S1.** Soil conditioning treatments have little impact on soil nutritional properties. **(A)** Soil organic carbon, **(B)** Bioavailable phosphorus, **(C)** Organic nitrogen, and **(D)** Organic matter content in non-conditioned control soils and soils conditioned with *Photorhabdus*-infected insect cadavers or mechanically killed larvae (MK). IDs 22–323 refer to the different *Photorhabdus* strains used for larval infection (Table 1). Box-and-whisker plots show minimum, first quartile, median, third quartile, and maximum values. n.s. not statistically significant relative to controls, determined by the Tukey's honest significant difference test, $p < 0.05$.

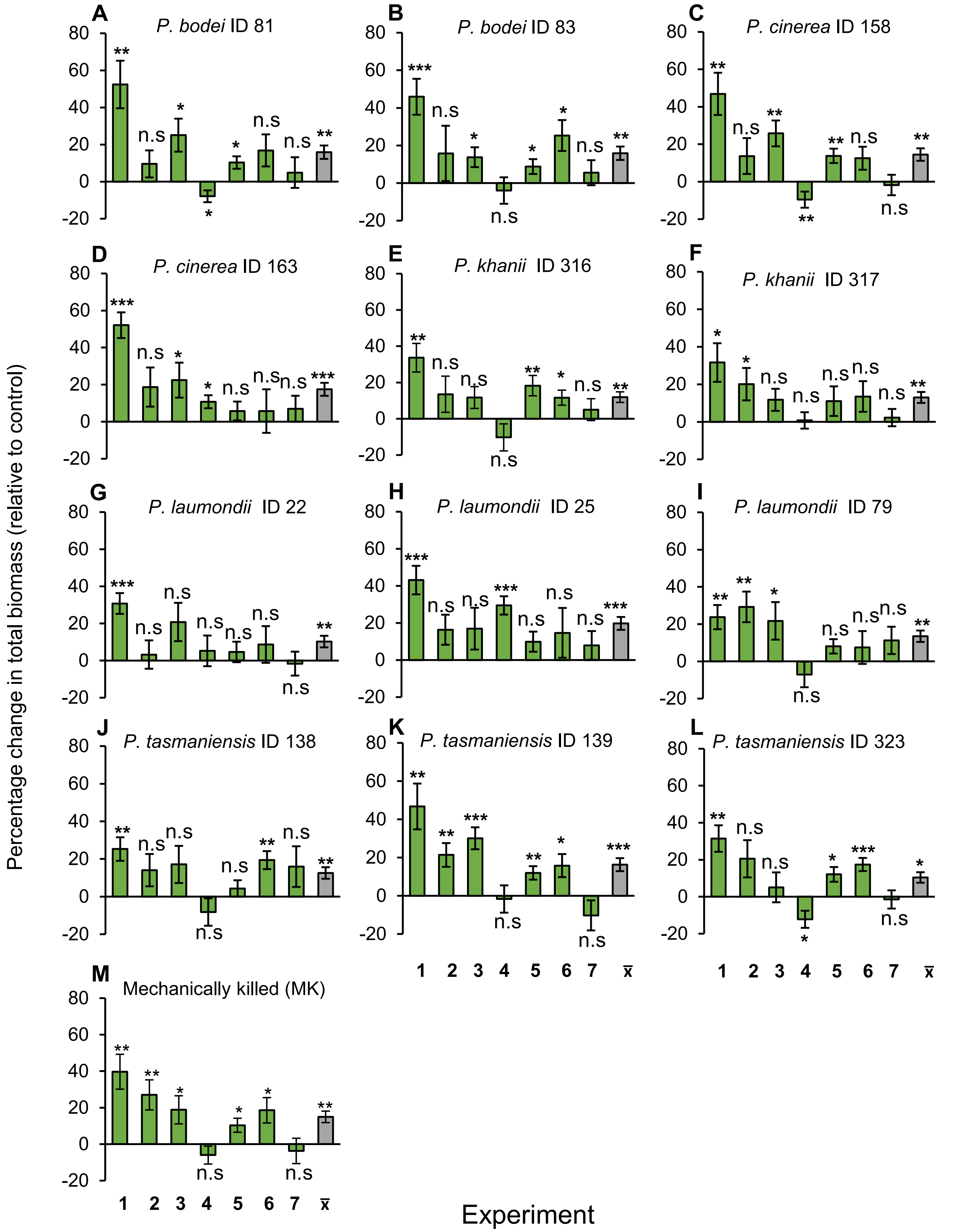


**Figure S2.** Plants often accumulate more biomass when grown on conditioned soils. **(A-L)** Percentage change (relative to controls) (± SE) in total biomass of plants grown in soil conditioned with *Photorhabdus*-infected insect cadavers, or **(M)** with mechanically killed larvae (MK). These experiments were conducted seven independent times with 10 replicates each time. The green bars show the percentage change in each experiment and the grey bars show the mean ($\bar{x}$) of all experiments. Asterisks above bars indicate significant increases in plant biomass (*: $p < 0.05$, **: $p < 0.01$, ***: $p < 0.001$; one-sample *t*-test). n.s. not statistically significant.

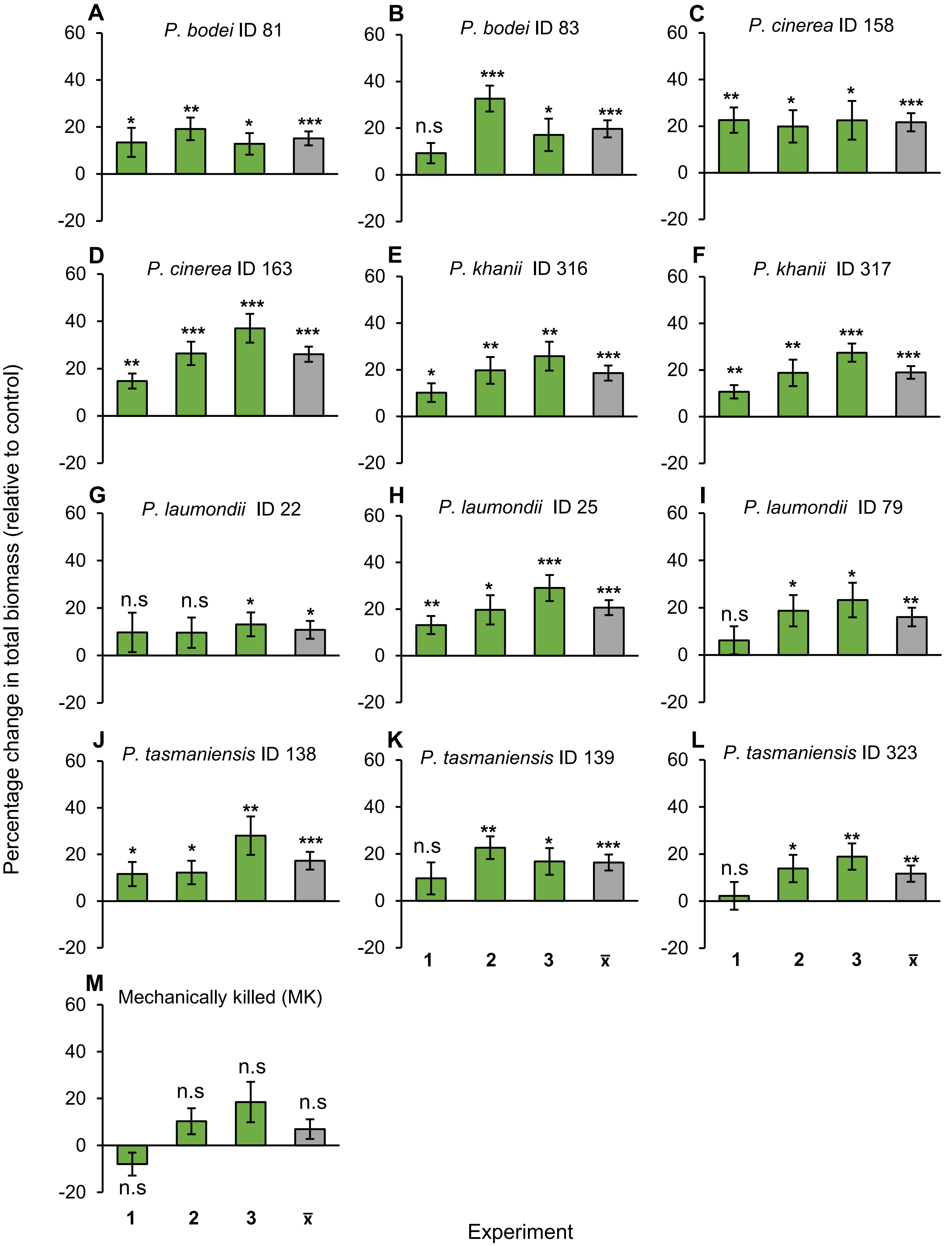


**Figure S3.** Plants often accumulate more biomass when grown on conditioned soil. **(A-L)** Percentage change (relative to controls) (± SE) in total biomass of plants grown in soil conditioned with aqueous extracts of *Photorhabdus*-infected insect cadavers, or **(M)** with aqueous extracts of mechanically killed larvae (MK). These experiments were conducted three independent times with 10 replicates each time. The green bars show the percentage change in each experiment and the grey bars show the mean (x̄) of all experiments. Asterisks above bars indicate significant increases in plant biomass (*: $p$ < 0.05, **: $p$ < 0.01, ***: p<0.001; one-sample $t$-test). n.s. not statistically significant.

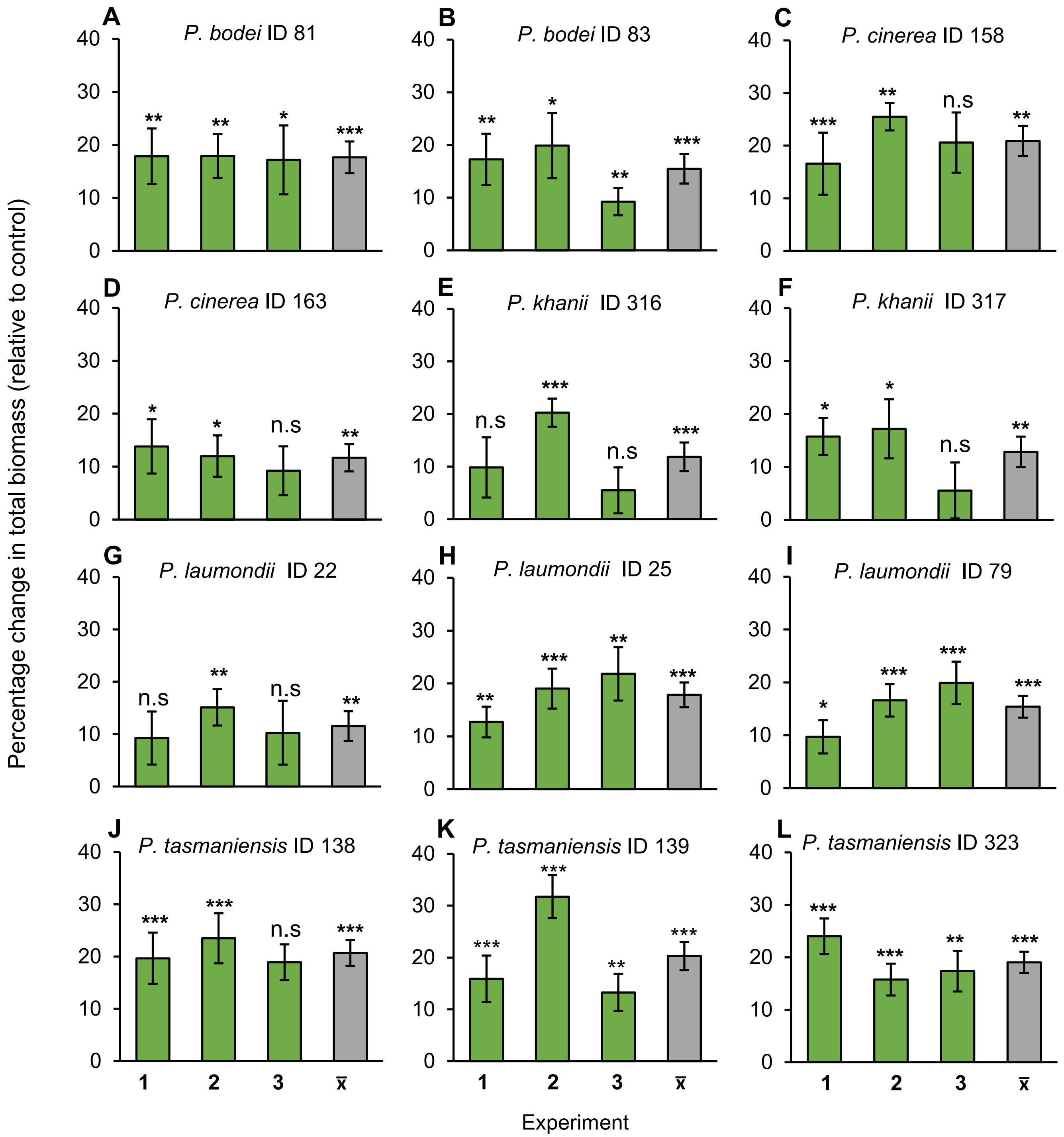


**Figure S4.** Plants accumulate more biomass when grown on conditioned soils. **(A-L)** Percentage change (relative to controls) (± SE) in total biomass of plants grown in soil conditioned with *Photorhabdus* cell-free supernatants. These experiments were conducted three independent times with 10 replicates each time. The green bars represent the total biomass accumulated in each experiment, and the grey bars show the mean (x̄) total biomass of all experiments. Asterisks above bars indicate significant increases in plant biomass (*: $p < 0.05$, **: $p < 0.01$, ***: $p < 0.001$; one-sample *t*-test). n.s. not statistically significant.

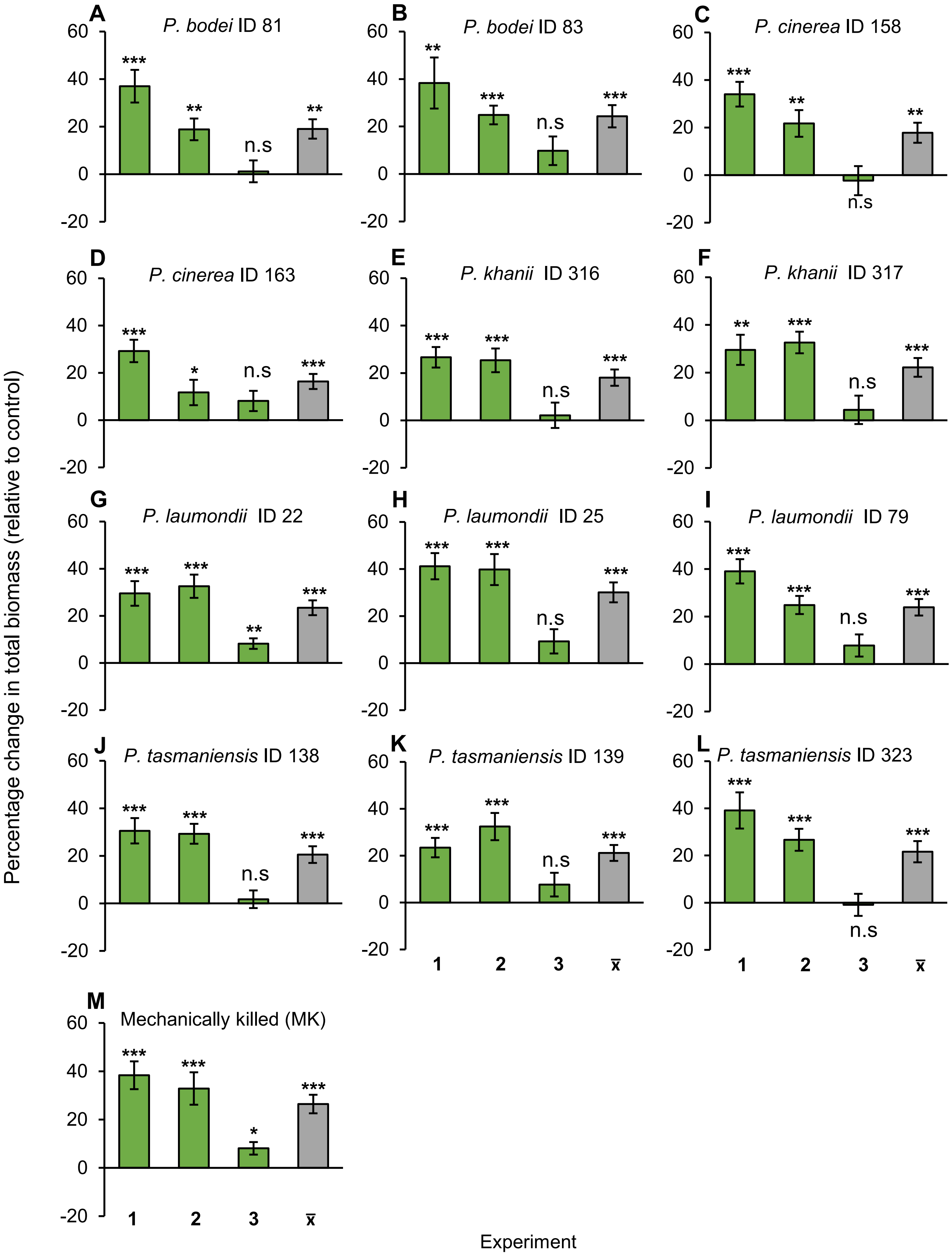


**Figure S5.** Plants often accumulate more biomass when grown on complemented soils. **(A-L)** Percentage change (relative to controls) (± SE) in total biomass of plants grown in autoclaved soils complemented with soil previously conditioned with *Photorhabdus*-infected insect cadavers or **(M)** mechanically killed larvae (MK). These experiments were conducted three independent times with 10 replicates each time. The green bars represent the total biomass accumulated in each experiment, and the grey bars show the mean ($\bar{x}$) total biomass of all experiments. Asterisks above bars indicate significant increases in plant biomass (*: $p < 0.05$, **: $p < 0.01$, ***: $p < 0.001$; one-sample $t$-test). n.s. not statistically significant.

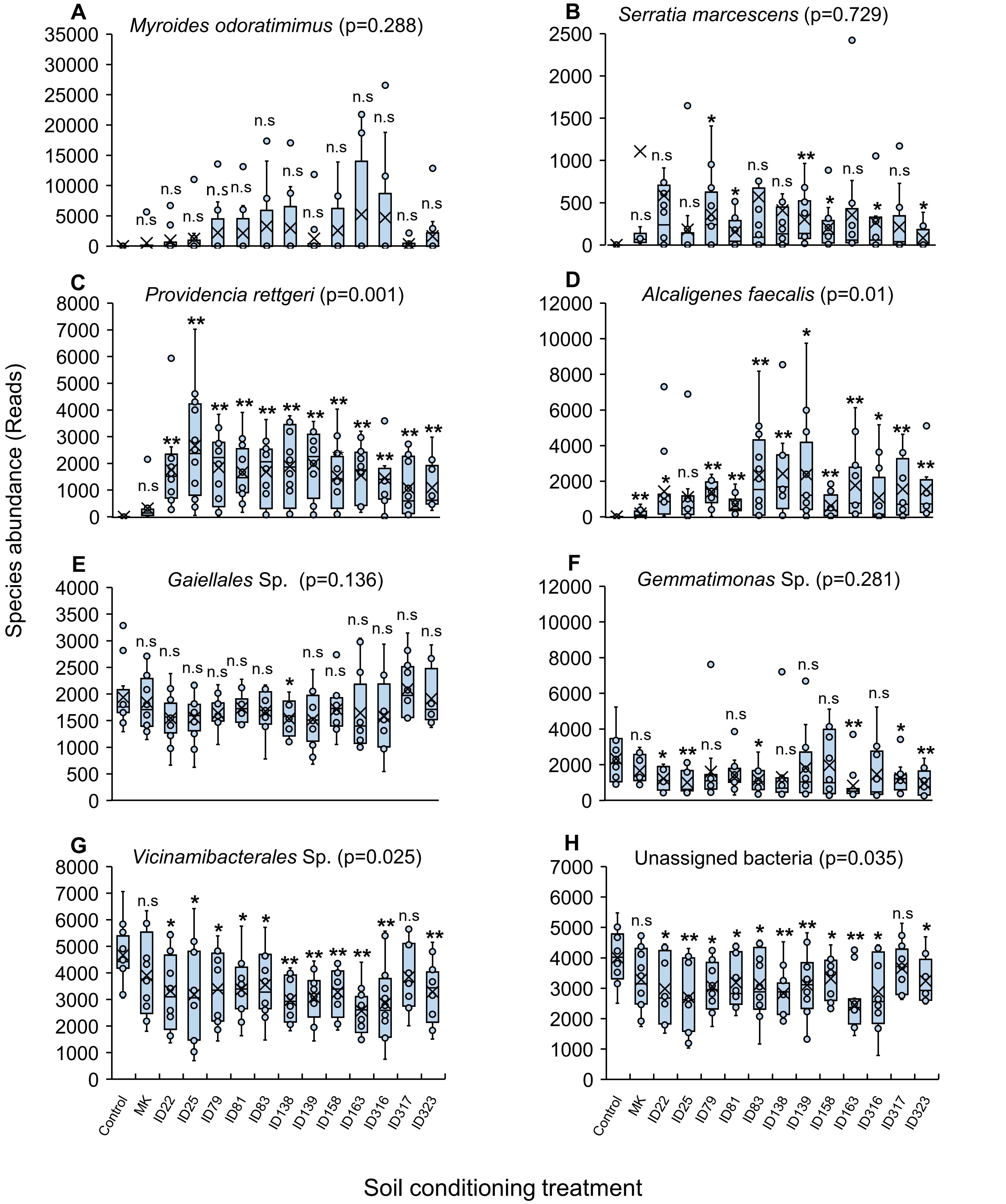


**Figure S6.** Soil conditioning treatments restructure soil bacterial communities. (**A–H**) Abundance of bacterial species in non-conditioned control soils and in soils conditioned with mechanically killed larvae (MK) or with *Photorhabdus*-infected insect cadavers. IDs 22–323 refer to the different *Photorhabdus* strains used for larval infections (Table 1). Boxes represent the interquartile range ($25^{th}$ – $75^{th}$ percentiles). In the boxplots, dots represent raw data, black crosses (X) indicate the median values of all variables, and whiskers indicate the minimum and maximum values. Asterisks above the bars indicate significant differences relative to the control ($p < 0.05$), determined by one-way ANOVA followed by Tukey's HSD test. n.s. not statistically significant.

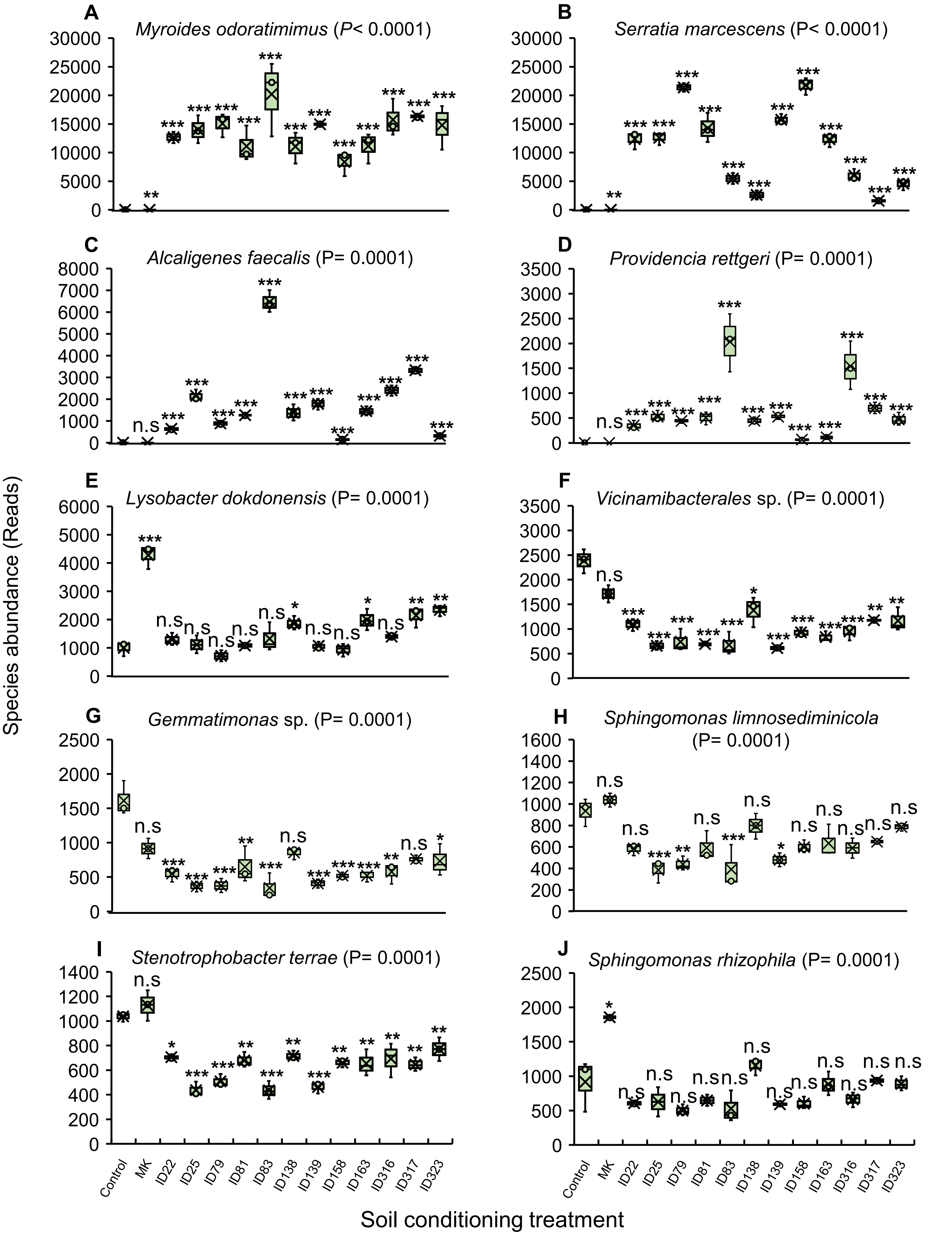


**Figure S7.** Soil conditioning treatments restructure the soil bacterial communities. (**A–J**) Abundance of bacterial species in non-conditioned control soils and in soils conditioned with aqueous extracts of mechanically killed larvae (MK) or with aqueous extracts of *Photorhabdus*-infected insect cadavers. IDs 22–323 refer to the different *Photorhabdus* strains used for larval infections (Table 1). Boxes represent the interquartile range (25th – 75th percentiles). In the boxplots, dots represent raw data, black crosses (X) indicate the median values of all variables, and whiskers indicate the minimum and maximum values. Asterisks above the bars indicate significant differences relative to the control ($p$ < 0.05), determined by one-way ANOVA followed by Tukey's HSD test. n.s. not statistically significant.

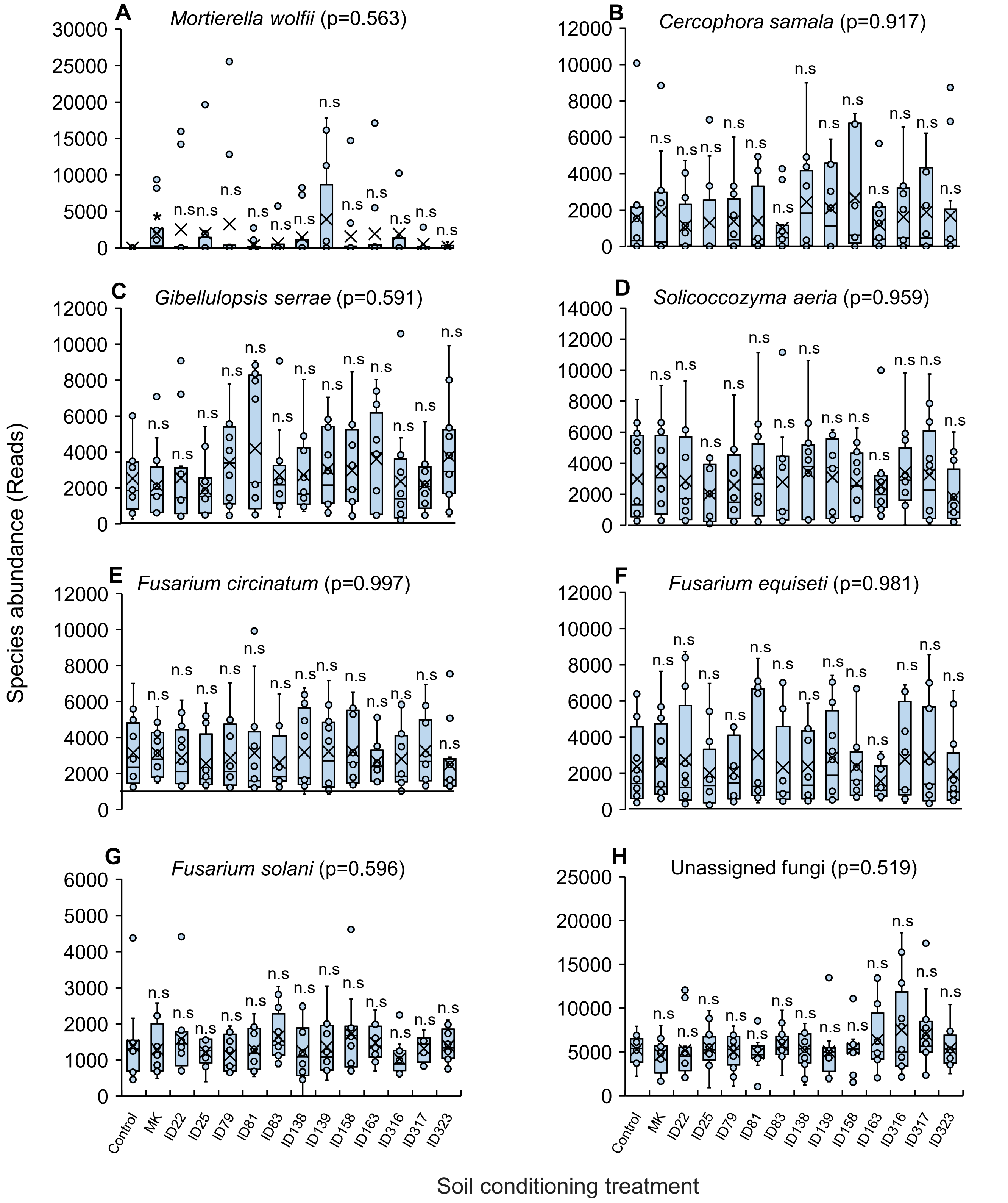


**Figure S8.** Soil conditioning treatments have little effect on the soil fungal communities. (**A–H**) Abundance of fungal species in non-conditioned control soils and in soils conditioned with mechanically killed larvae (MK) or with *Photorhabdus*-infected insect cadavers. IDs 22–323 refer to the different *Photorhabdus* strains used for larval infections (Table 1). Boxes represent the interquartile range ($25^{th}$ – $75^{th}$ percentiles). In the boxplots, dots represent raw data, black crosses (X) indicate the median values of all variables, and whiskers indicate the minimum and maximum values. Asterisks above the bars indicate significant differences relative to the control ($p$ < 0.05), determined by one-way ANOVA followed by Tukey's HSD test. n.s. not statistically significant.

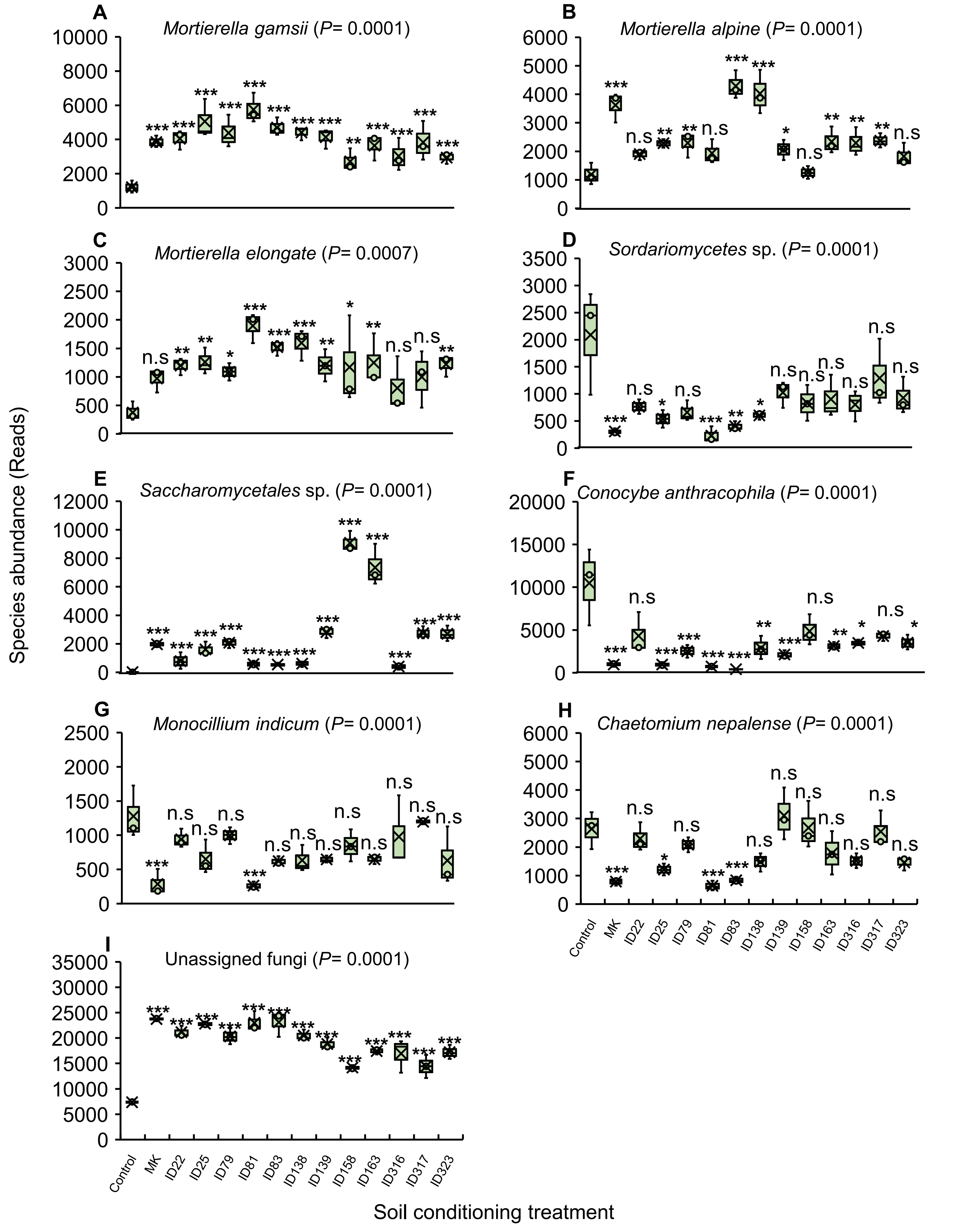


**Figure S9.** Soil conditioning treatments restructure the soil fungal communities. (**A–I)** Abundance of fungal species in non-conditioned control soils and in soils conditioned with aqueous extracts of mechanically killed larvae (MK) or with aqueous extracts of *Photorhabdus*-infected insect cadavers. IDs 22–323 refer to the different *Photorhabdus* strains used for larval infections (Table 1). Boxes represent the interquartile range (25th – 75th percentiles). In the boxplots, dots represent raw data, black crosses (X) indicate the median values of all variables, and whiskers indicate the minimum and maximum values. Asterisks above the bars indicate significant differences relative to the control ($p < 0.05$), determined by one-way ANOVA followed by Tukey's HSD test. n.s. not statistically significant.

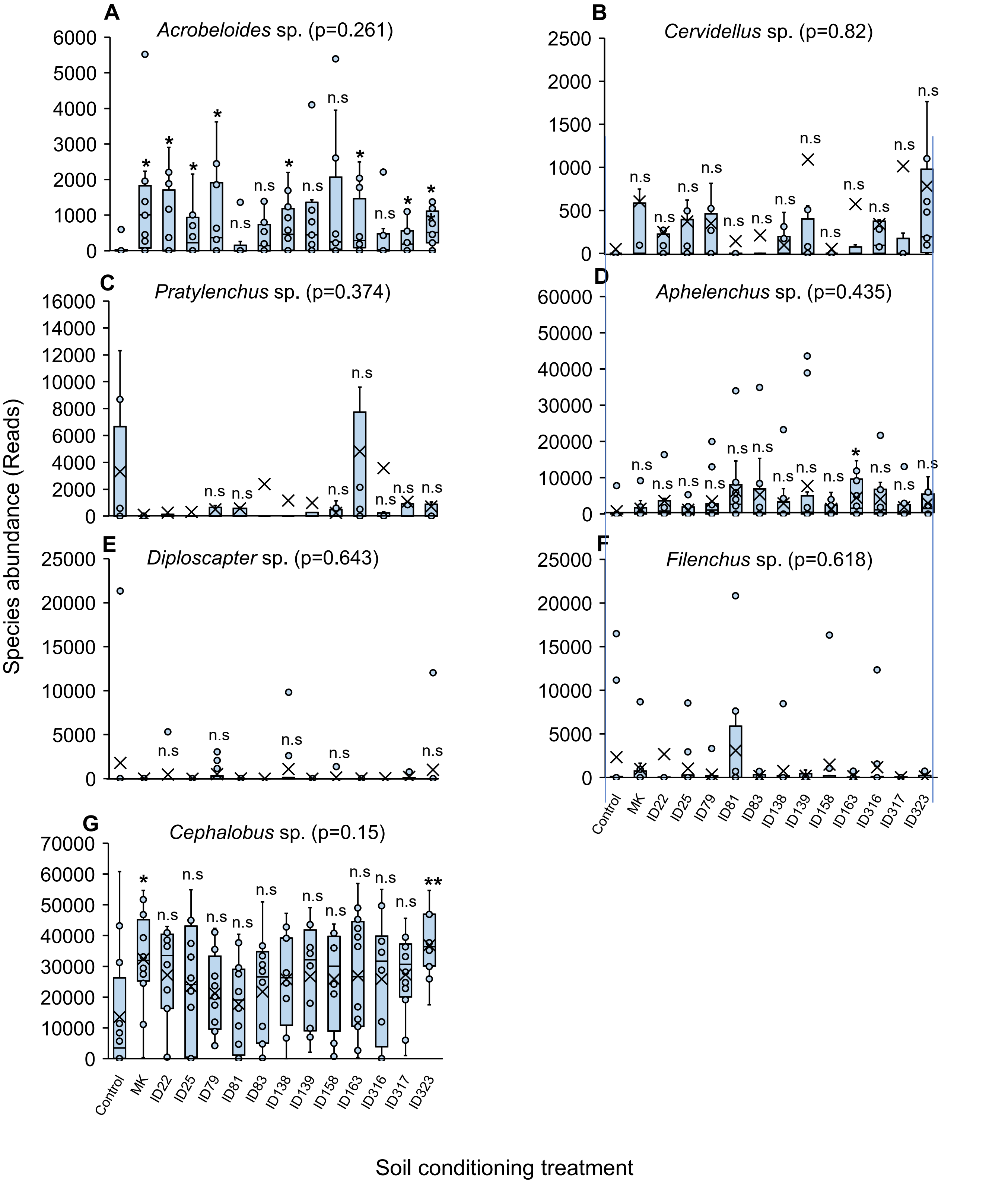


**Figure S10.** Soil conditioning treatments restructure the nematode communities. (**A–G**) Abundance of nematode species in non-conditioned control soils and in soils conditioned with mechanically killed larvae (MK) or with *Photorhabdus*-infected insect cadavers. IDs 22–323 refer to the different *Photorhabdus* strains used for larval infections (Table 1). Boxes represent the interquartile range ($25^{th}$ – $75^{th}$ percentiles). In the boxplots, dots represent raw data, black crosses (X) indicate the median values of all variables, and whiskers indicate the minimum and maximum values. Asterisks above the bars indicate significant differences relative to the control ($p < 0.05$), determined by one-way ANOVA followed by Tukey's HSD test. n.s. not statistically significant.

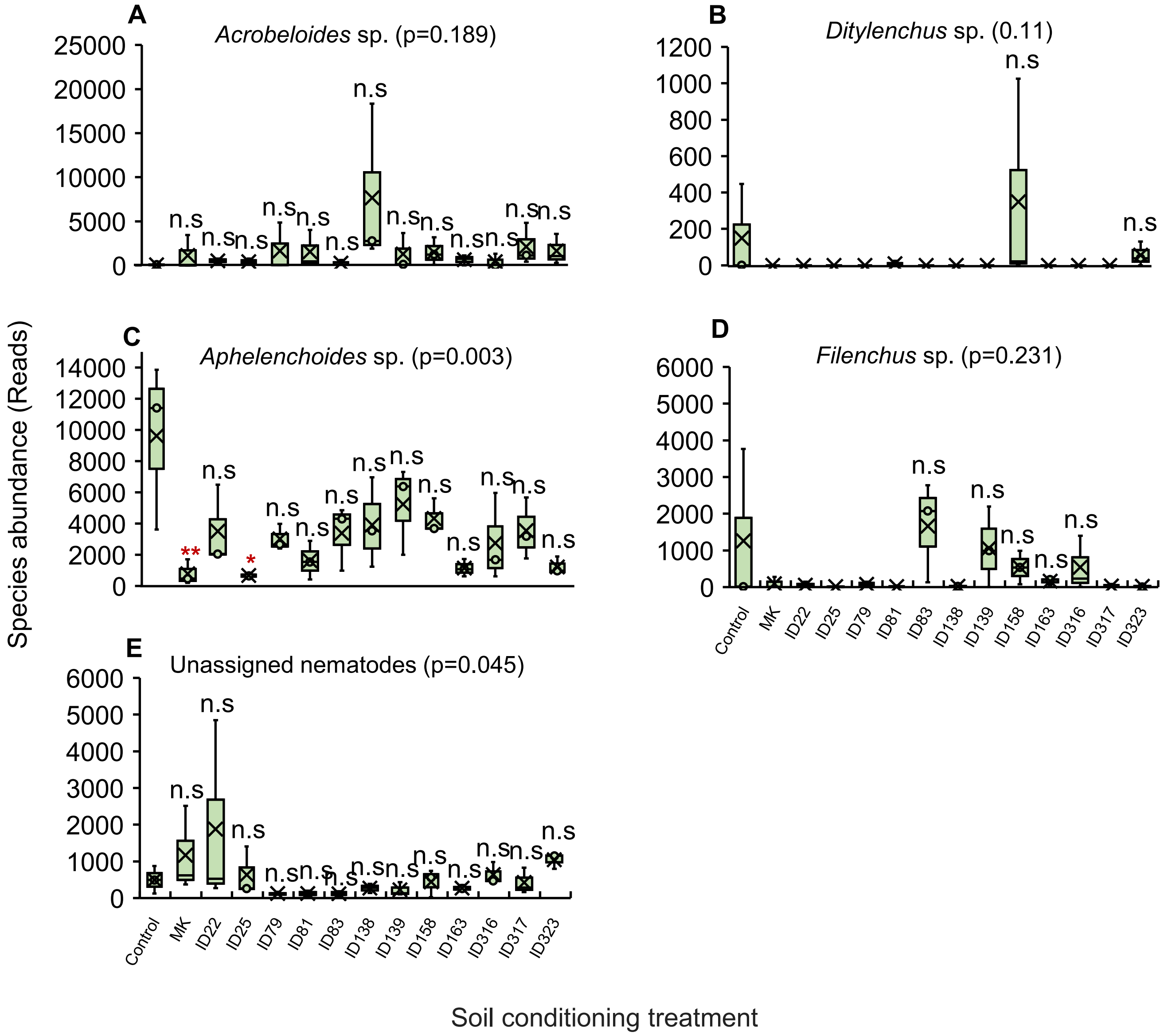


**Figure S11.** Soil conditioning treatments restructure the nematode communities. (**A–E)** Abundance of nematode species in non-conditioned control soils and in soils conditioned with aqueous extracts of mechanically killed larvae (MK) or with aqueous extracts of *Photorhabdus*-infected insect cadavers. IDs 22–323 refer to the different *Photorhabdus* strains used for larval infections (Table 1). Boxes represent the interquartile range ($25^{th}$ – $75^{th}$ percentiles). In the boxplots, dots represent raw data, black crosses (X) indicate the median values of all variables, and whiskers indicate the minimum and maximum values. Asterisks above the bars indicate significant differences relative to the control ($p < 0.05$), determined by one-way ANOVA followed by Tukey's HSD test. n.s. not statistically significant.

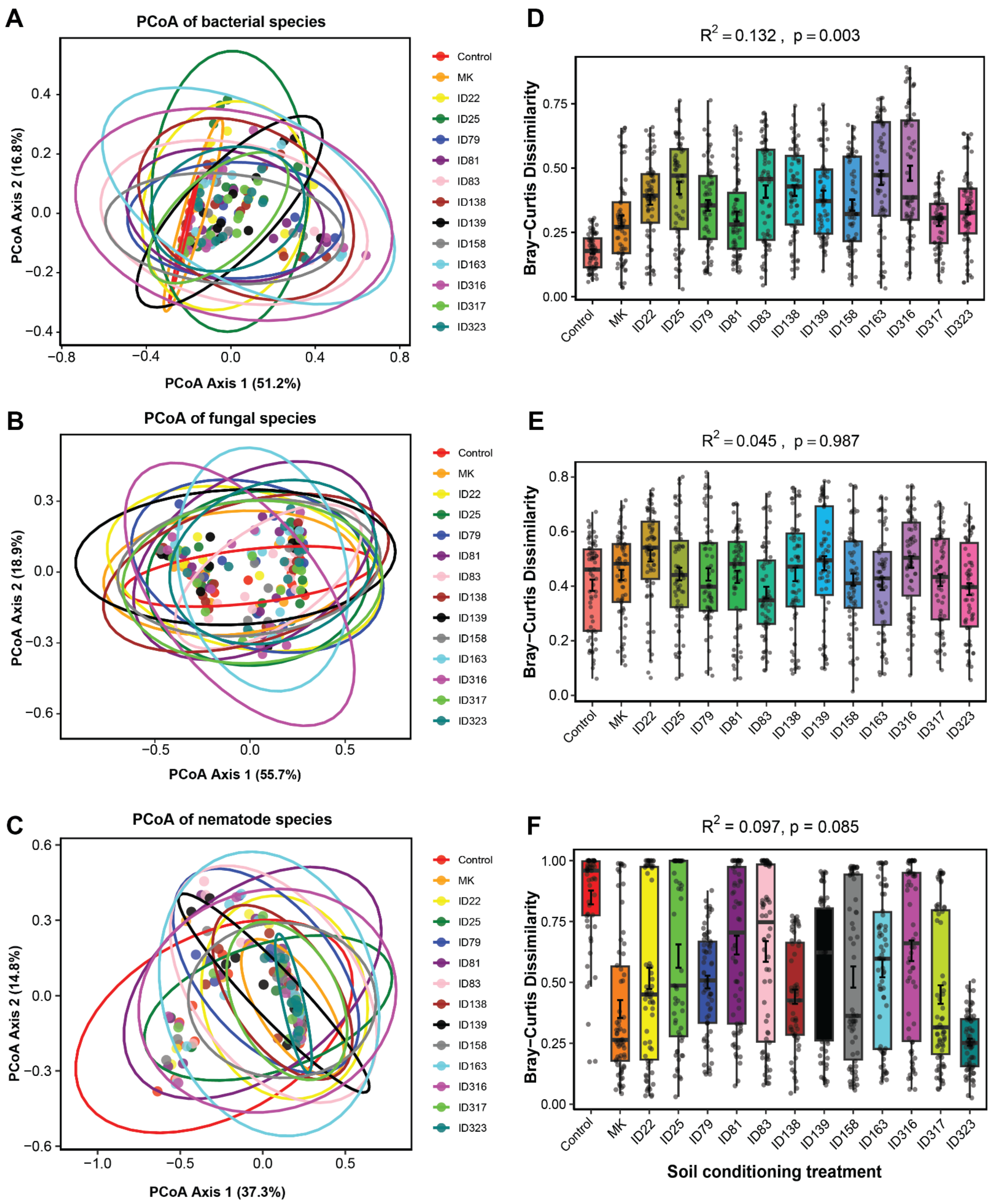


**Figure S12.** Soil conditioning treatments alter bacterial and fungal communities, and to a lesser extent, nematode communities. **(A–C)** Principal coordinates analyses (PCoA) based on Bray–Curtis dissimilarities and **(D–F)** PERMANOVA results for bacterial, fungal and nematode communities in non-conditioned soil and in soil conditioned with aqueous extracts of mechanically killed larvae (MK) or aqueous extracts of *Photorhabdus*-infected insect cadavers. IDs 22–323 denote the *Photorhabdus* strains used for larval infection (Table 1). Ellipses represent 95% confidence intervals for each treatment. $R^2$ values and *p*-values indicate the effect of soil conditioning on bacterial, fungal and nematode community composition.

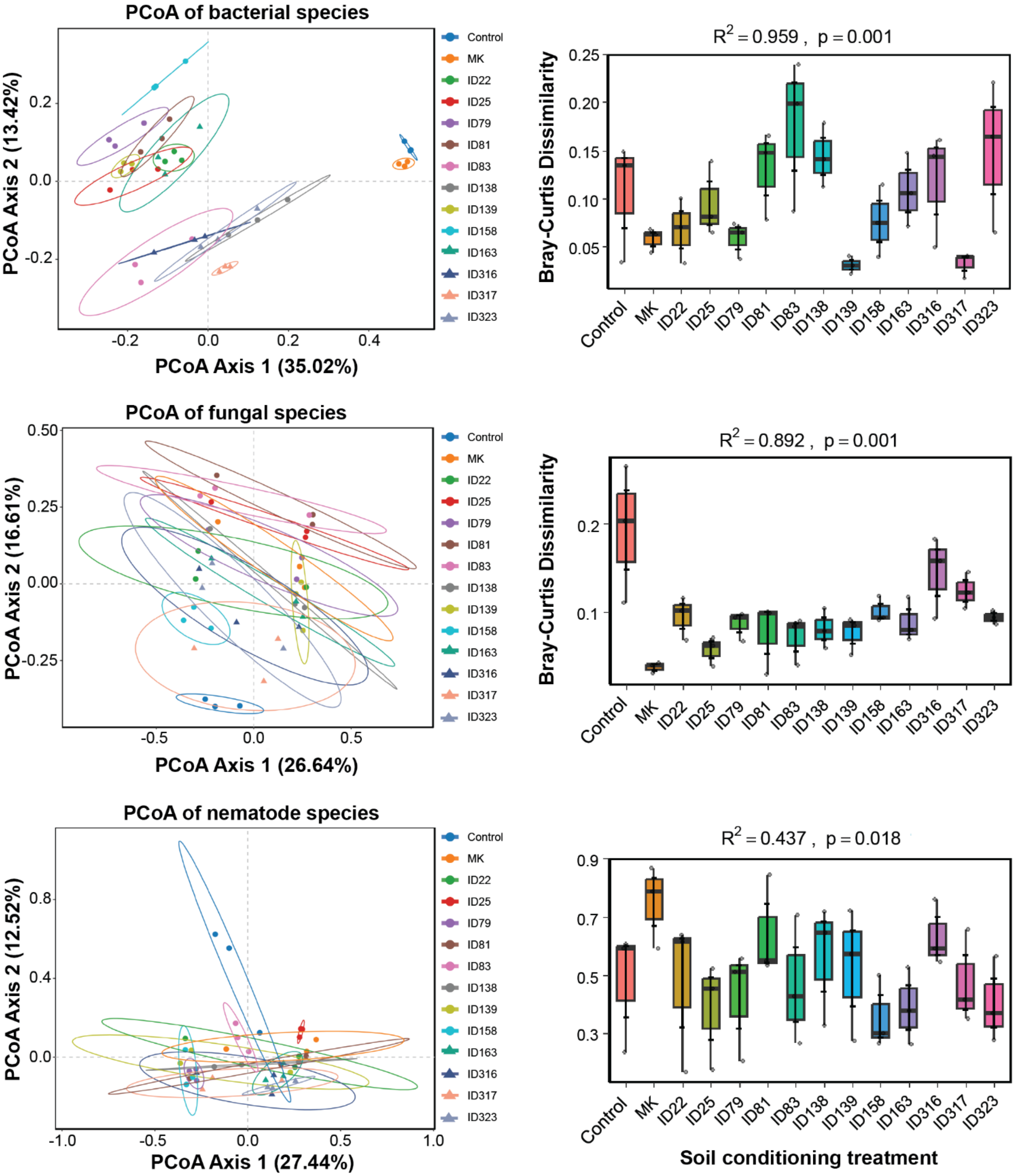


**Figure S13.** Soil conditioning treatments alter bacterial, fungal and nematode communities. **(A–C)** Principal coordinates analyses (PCoA) based on Bray–Curtis dissimilarities and **(D–F)** PERMANOVA results for bacterial, fungal and nematode communities in non-conditioned soil and in soil conditioned with aqueous extracts of mechanically killed larvae (MK) or with aqueous extracts of *Photorhabdus*-infected insect cadavers. IDs 22–323 denote the *Photorhabdus* strains used for larval infection (Table 1). Ellipses represent 95% confidence intervals for each treatment. $R^2$ values and *p*-values indicate the effect of soil conditioning on bacterial, fungal and nematode community composition.

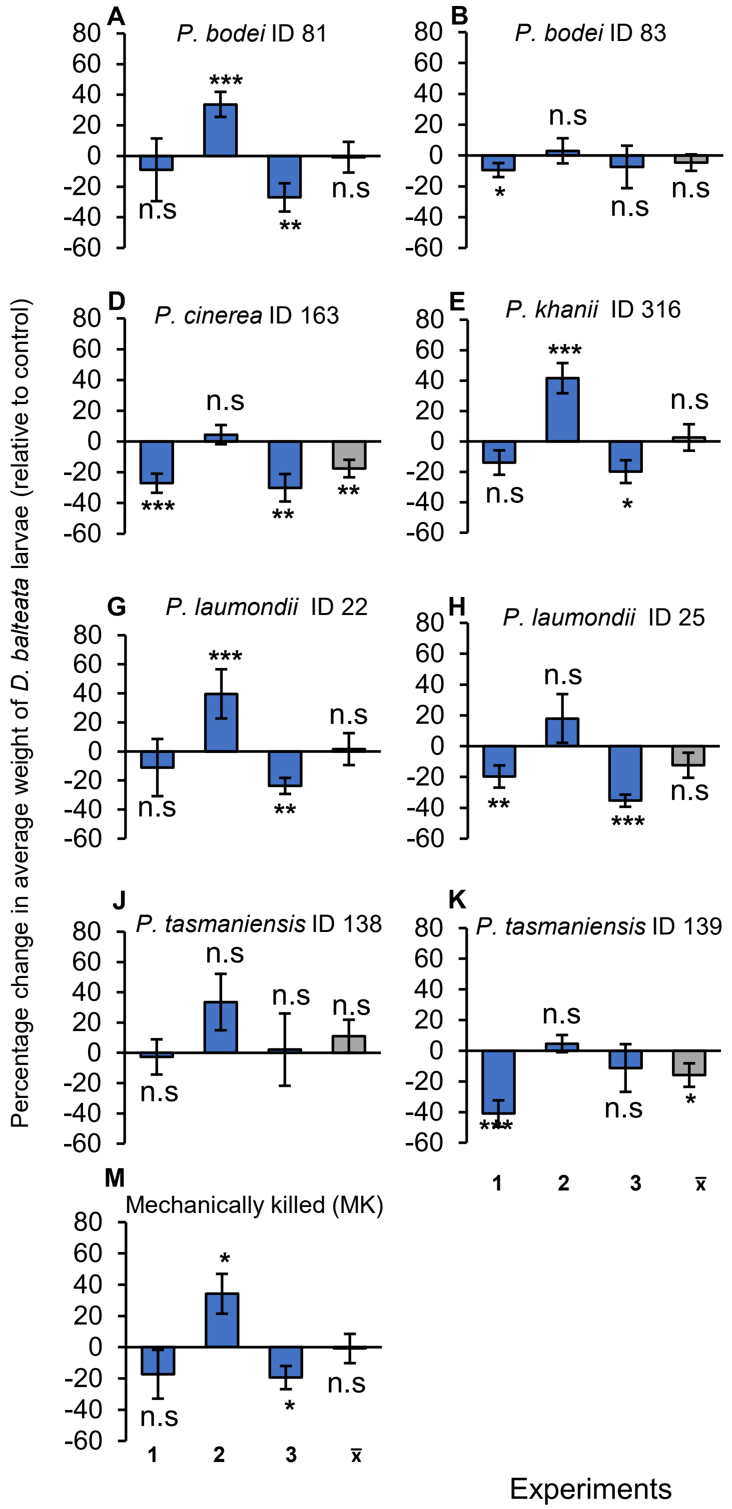


**Figure S14.** Plants grown on conditioned soils often resist the attack of *D. balteata*. **(A–M)** Percentage change (relative to controls) (± SE) in the performance of *D. balteata* on maize plants grown on soils conditioned with *Photorhabdus*-infected insect cadavers, or **(M)** mechanically killed larvae (MK). IDs 22–323 refer to the different *Photorhabdus* strains used for larval infections (Table 1). These experiments were conducted three independent times, with 5 replicates each time and 10 larvae per replicate. Asterisks above bars indicate significant changes in the average weight of *D. balteata* larvae (*: $p < 0.05$, **: $p < 0.01$, ***: $p < 0.001$; one-sample *t*-test). Positive values indicate weight gain and negative values indicate weight loss. n.s. not statistically significant.

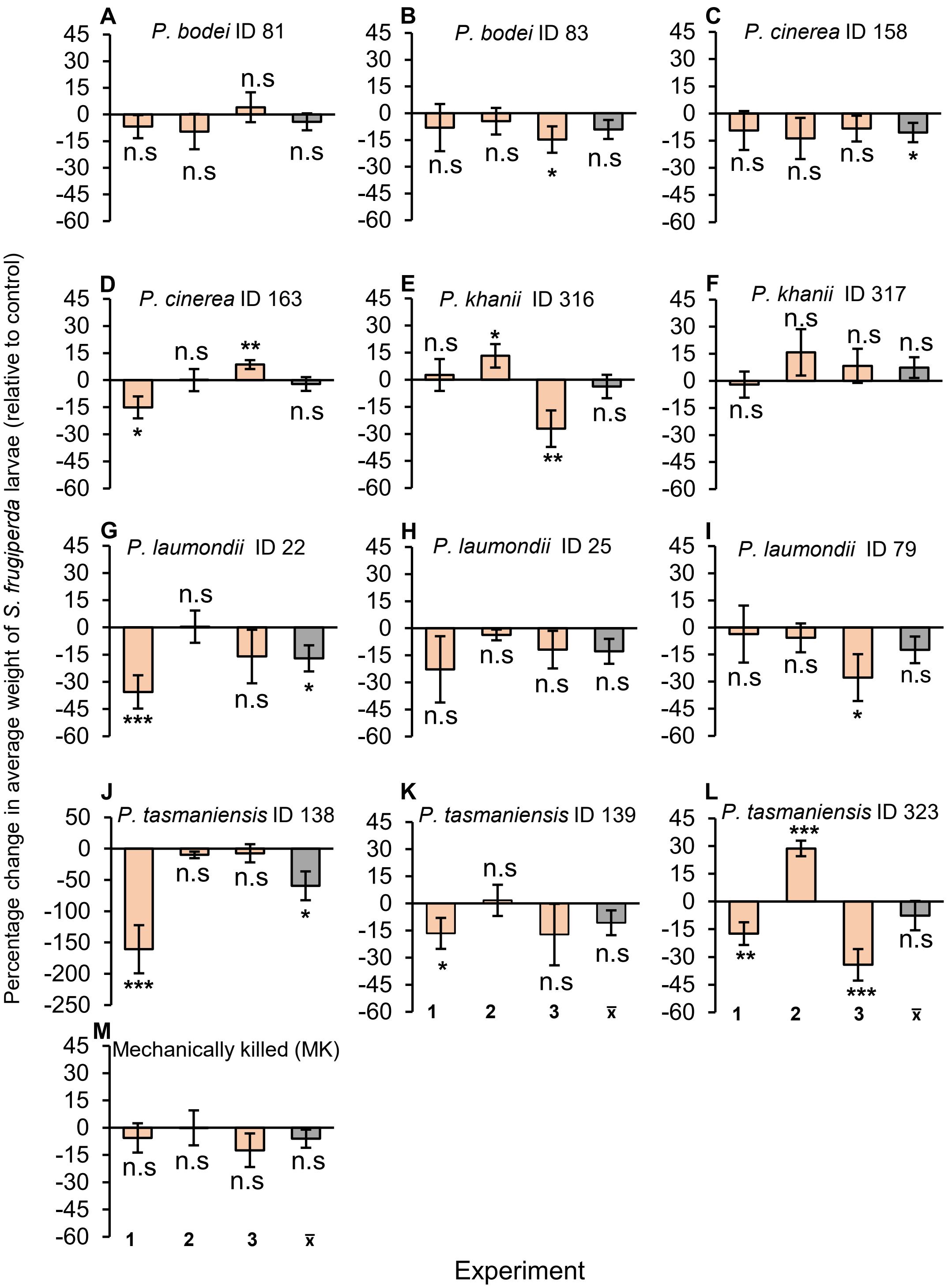


**Figure S15.** Plants grown on conditioned soils often resist the attack of *S. frugiperda*. **(A–M)** Percentage change (relative to controls) (± SE) in the performance of *S. frugiperda* on maize plants grown on soils conditioned with *Photorhabdus*-infected insect cadavers, or **(M)** mechanically killed larvae (MK). IDs 22–323 refer to the different *Photorhabdus* strains used for larval infections (Table 1). These experiments were conducted three independent times, with 5 replicates each time and 10 larvae per replicate. Asterisks above bars indicate significant reductions or increases in the average weight of *S. frugiperda* larvae (*: $p < 0.05$, **: $p < 0.01$, ***: $p < 0.001$; one-sample *t*-test). Positive values indicate weight gain and negative values indicate weight loss. n.s. not statistically significant.

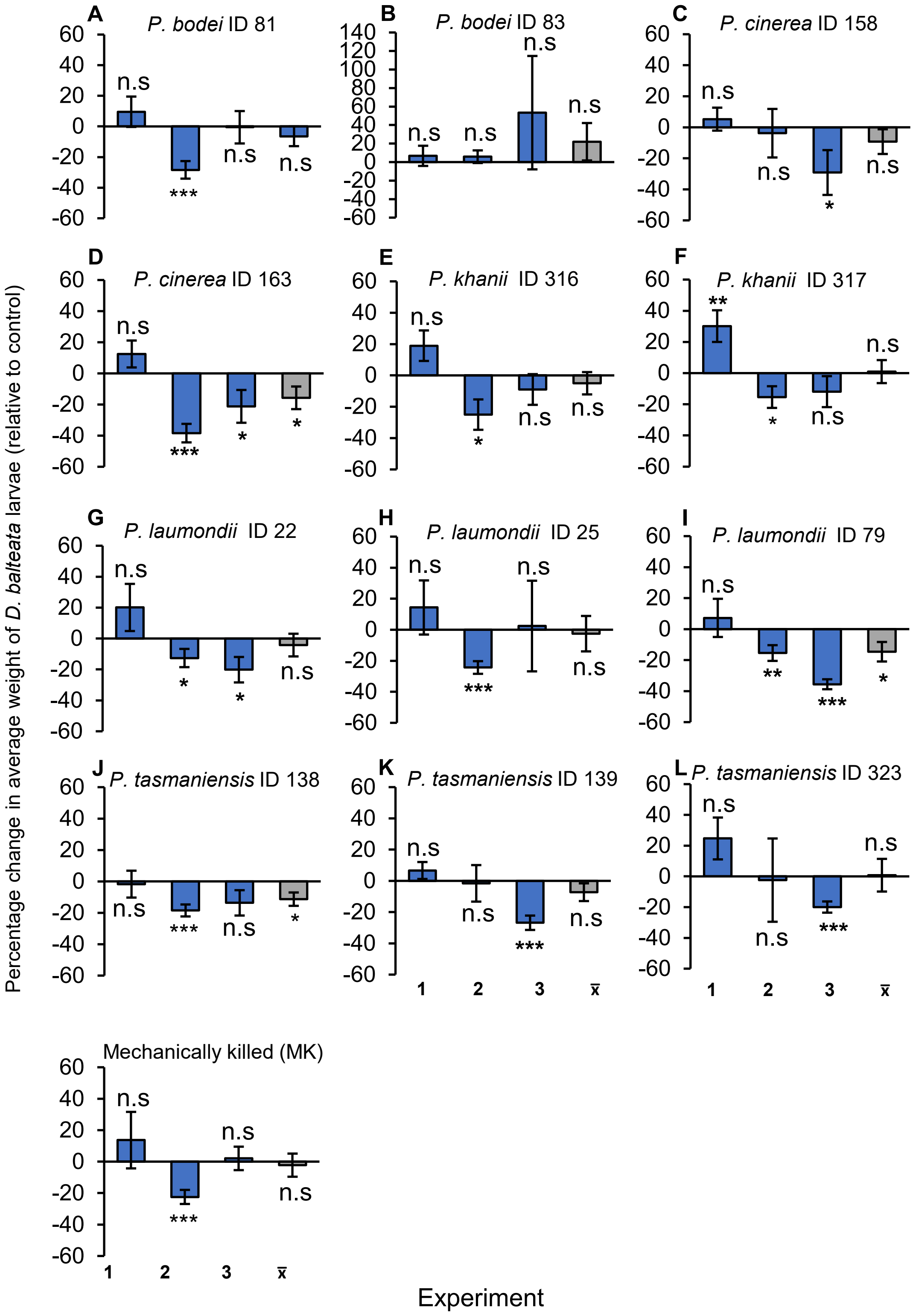


**Figure S16.** Plants grown on conditioned soils often resist the attack of *D. balteata*. **(A–M)** Percentage change (relative to controls) (± SE) in the performance of *D. balteata* on maize plants grown on soils conditioned with aqueous extracts of *Photorhabdus*-infected insect cadavers, or **(M)** with aqueous extracts of mechanically killed larvae (MK). IDs 22–323 refer to the different *Photorhabdus* strains used for larval infections (Table 1). These experiments were conducted three independent times, with 5 replicates each time and 10 larvae per replicate. Asterisks above bars indicate significant changes in the average weight of *D. balteata* larvae (*: $p < 0.05$, **: $p < 0.01$, ***: $p < 0.001$; one-sample *t*-test). Positive values indicate weight gain and negative values indicate weight loss. n.s. not statistically significant.

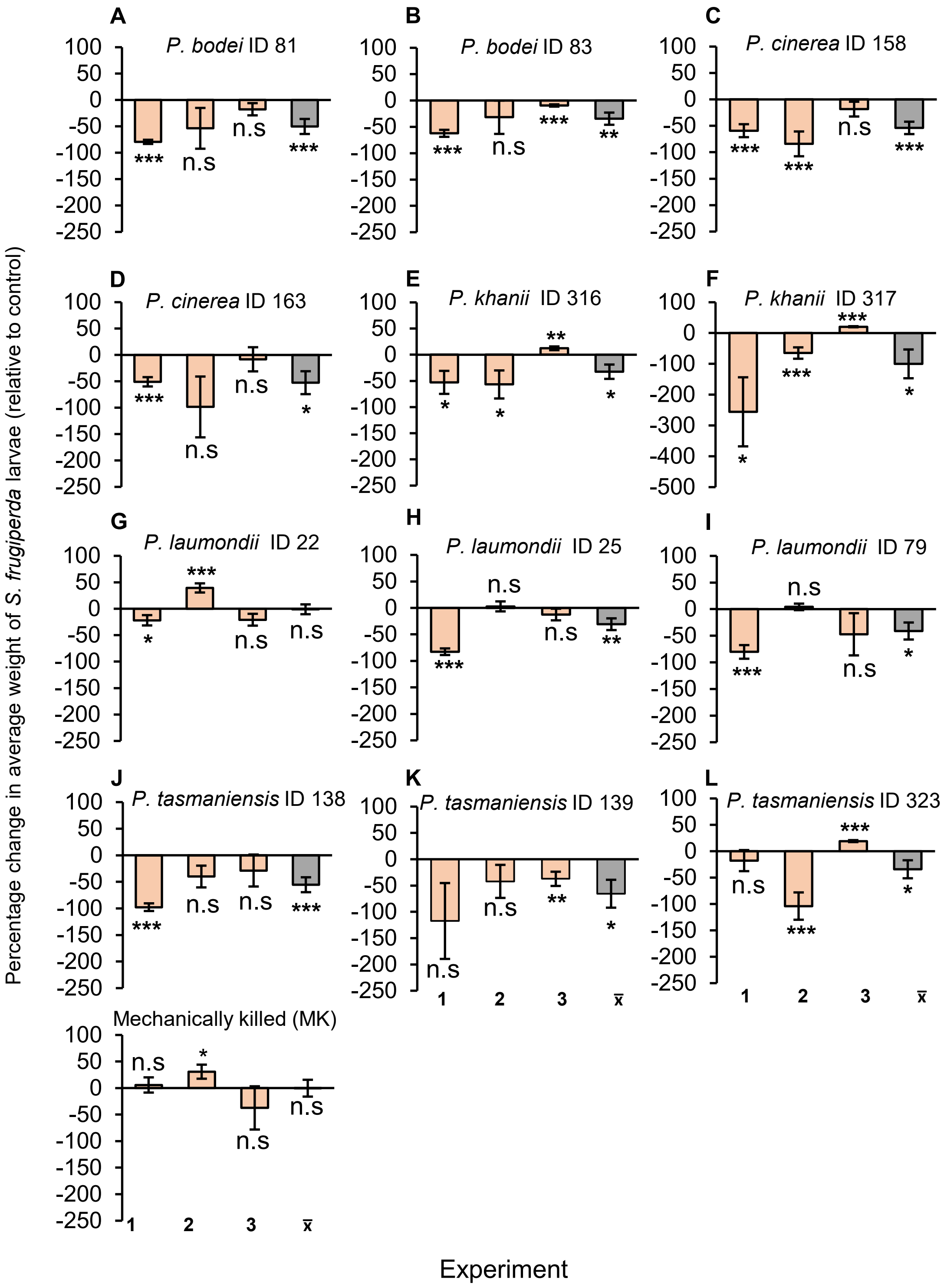


**Figure S17.** Plants grown on conditioned soils often resist the attack of *S. frugiperda*. **(A–M)** Percentage change (relative to controls) (± SE) in the performance of *S. frugiperda* on maize plants grown on soils conditioned with aqueous extracts of *Photorhabdus*-infected insect cadavers, or **(M)** with aqueous extracts of mechanically killed larvae (MK). IDs 22–323 refer to the different *Photorhabdus* strains used for larval infections (Table 1). These experiments were conducted three independent times, with 5 replicates each time and 10 larvae per replicate. Asterisks above bars indicate significant reductions or increases in the average weight of *S. frugiperda* larvae (*: $p < 0.05$, **: $p < 0.01$, ***: $p < 0.001$; one-sample *t*-test). Positive values indicate weight gain and negative values indicate weight loss. n.s. not statistically significant.

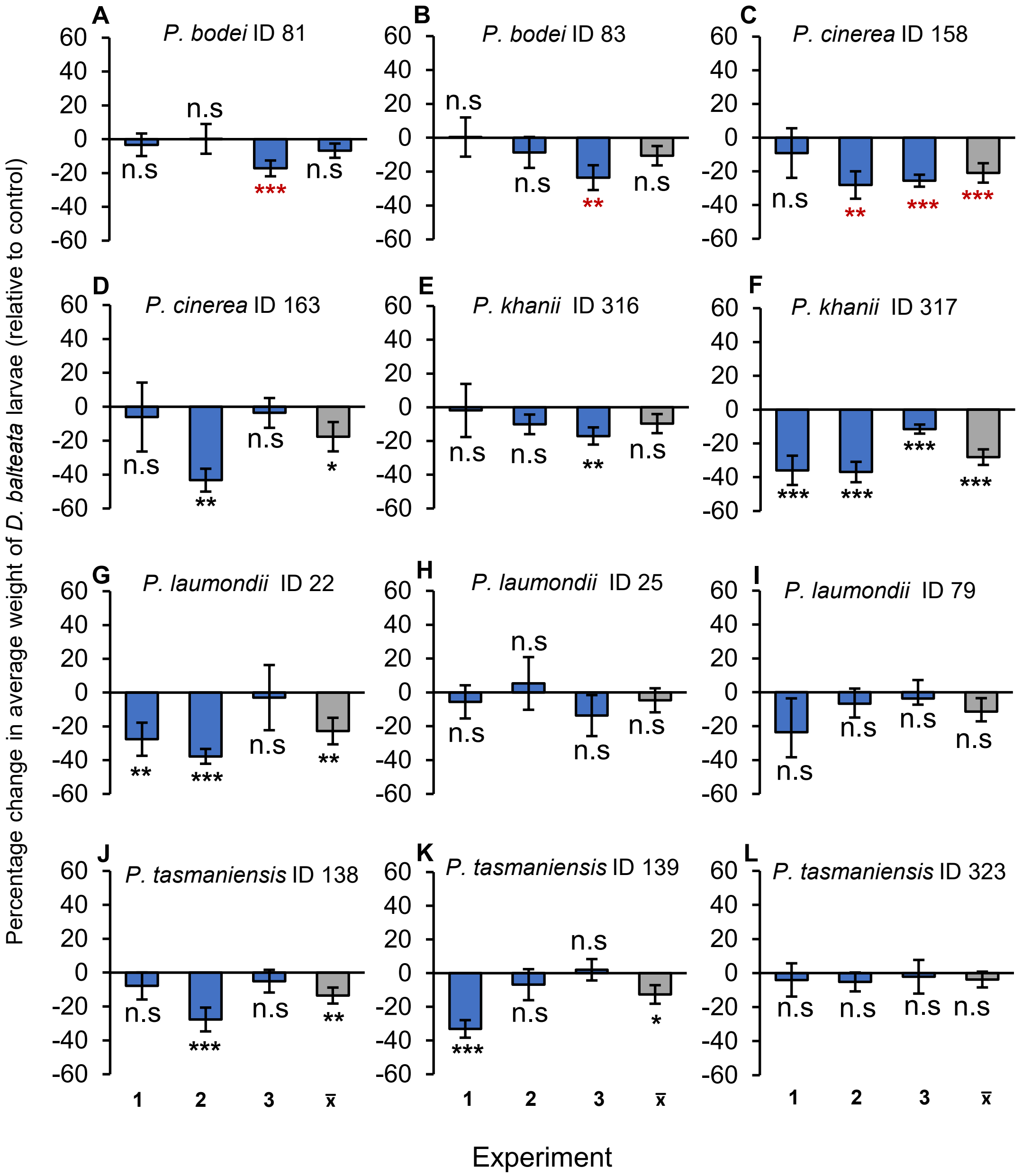


**Figure S18.** Plants grown on conditioned soils often resist the attack of *D. balteata*. **(A–L)** Percentage change (relative to controls) (± SE) in the performance of *D. balteata* on maize plants grown on soils conditioned with *Photorhabdus* cell-free supernatants. IDs 22–323 refer to the different *Photorhabdus* strains used for larval infections (Table 1). These experiments were conducted three independent times, with 5 replicates each time and 10 larvae per replicate. Asterisks above bars indicate significant changes in the average weight of *D. balteata* larvae (*: $p < 0.05$, **: $p < 0.01$, ***: $p < 0.001$; one-sample $t$-test). Positive values indicate weight gain and negative values indicate weight loss. n.s. not statistically significant.

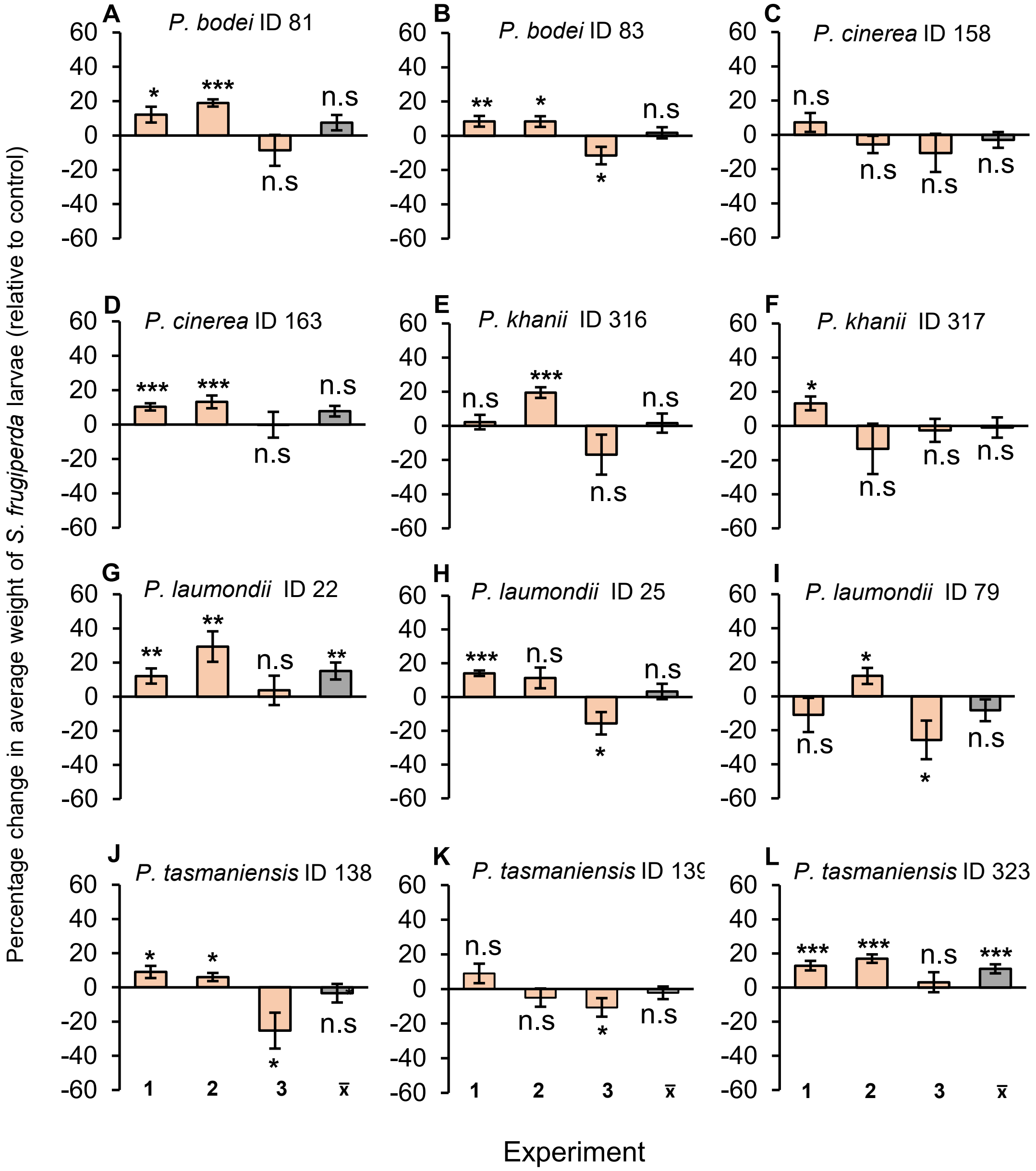


**Figure S19.** Plants grown on conditioned soils often resist the attack of *S. frugiperda*. **(A–M)** Percentage change (relative to controls) (± SE) in the performance of *S. frugiperda* on maize plants grown on soils conditioned with *Photorhabdus* cell-free supernatants. IDs 22–323 refer to the different *Photorhabdus* strains used for larval infections (Table 1). These experiments were conducted three independent times, with 5 replicates each time and 10 larvae per replicate. Asterisks above bars indicate significant reductions or increases in the average weight of *S. frugiperda* larvae (*: $p < 0.05$, **: $p < 0.01$, ***: $p < 0.001$; one-sample *t*-test). Positive values indicate weight gain and negative values indicate weight loss. n.s. not statistically significant.

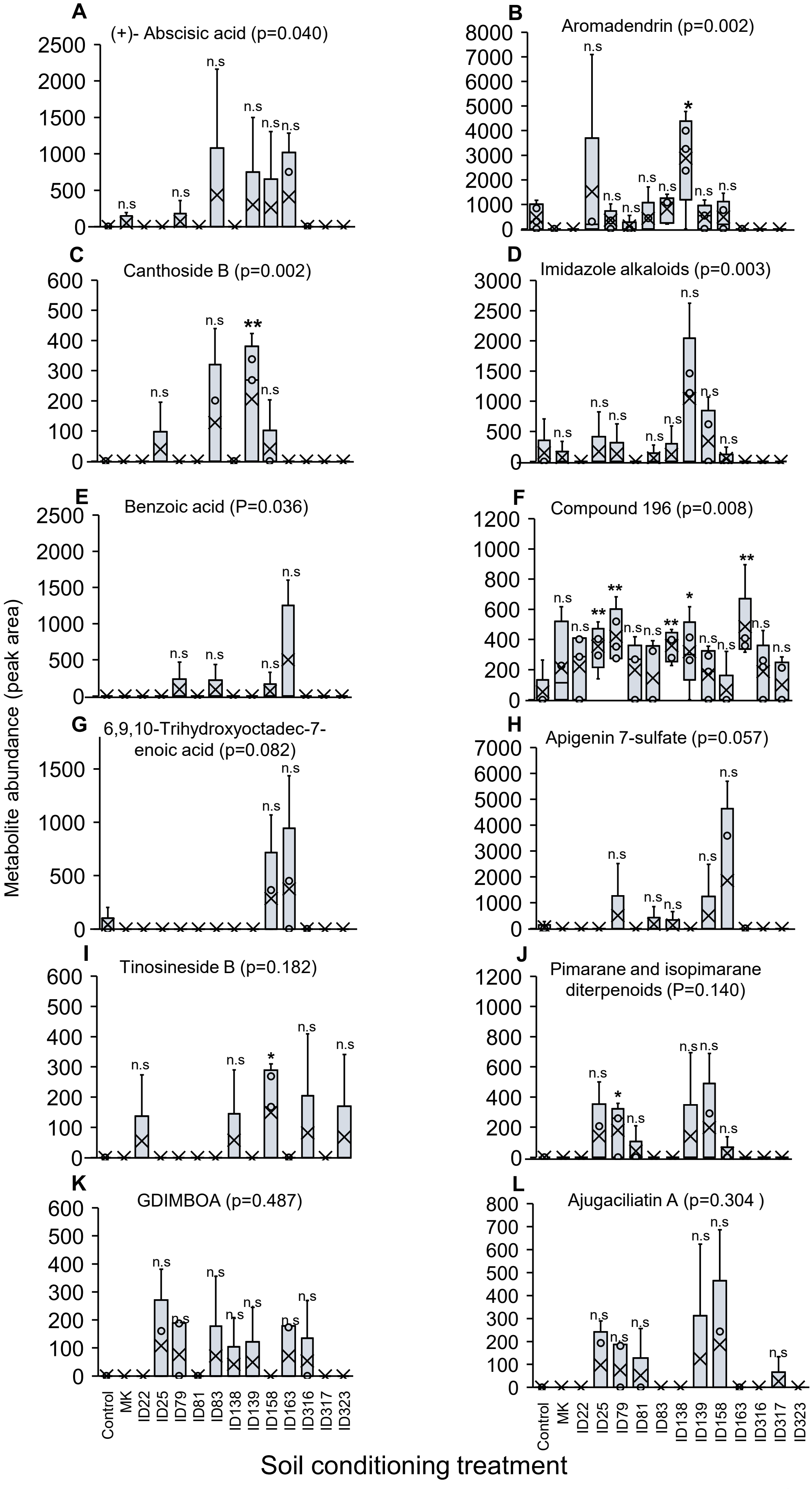


**Figure S20.** Plant roots respond at the metabolic level to soil conditioning treatments. (**A–L)** Abundance of upregulated low molecular weight metabolites in the roots of plants grown on non-conditioned control soil and in soils conditioned with mechanically killed larvae (MK) or with *Photorhabdus*-infected insect cadavers. IDs 22–323 refer to the different *Photorhabdus* strains used for larval infections (Table 1). Boxes represent the interquartile range (25$^{th}$ – 75$^{th}$ percentiles). In the boxplots, dots represent the raw data, black crosses (X) indicate the median values of all variables, and whiskers indicate the minimum and maximum values. Asterisks above the bars indicate significant differences relative to the control ($p$ < 0.05), assessed by one-way ANOVA followed by Tukey's HSD test. n.s. not statistically significant.

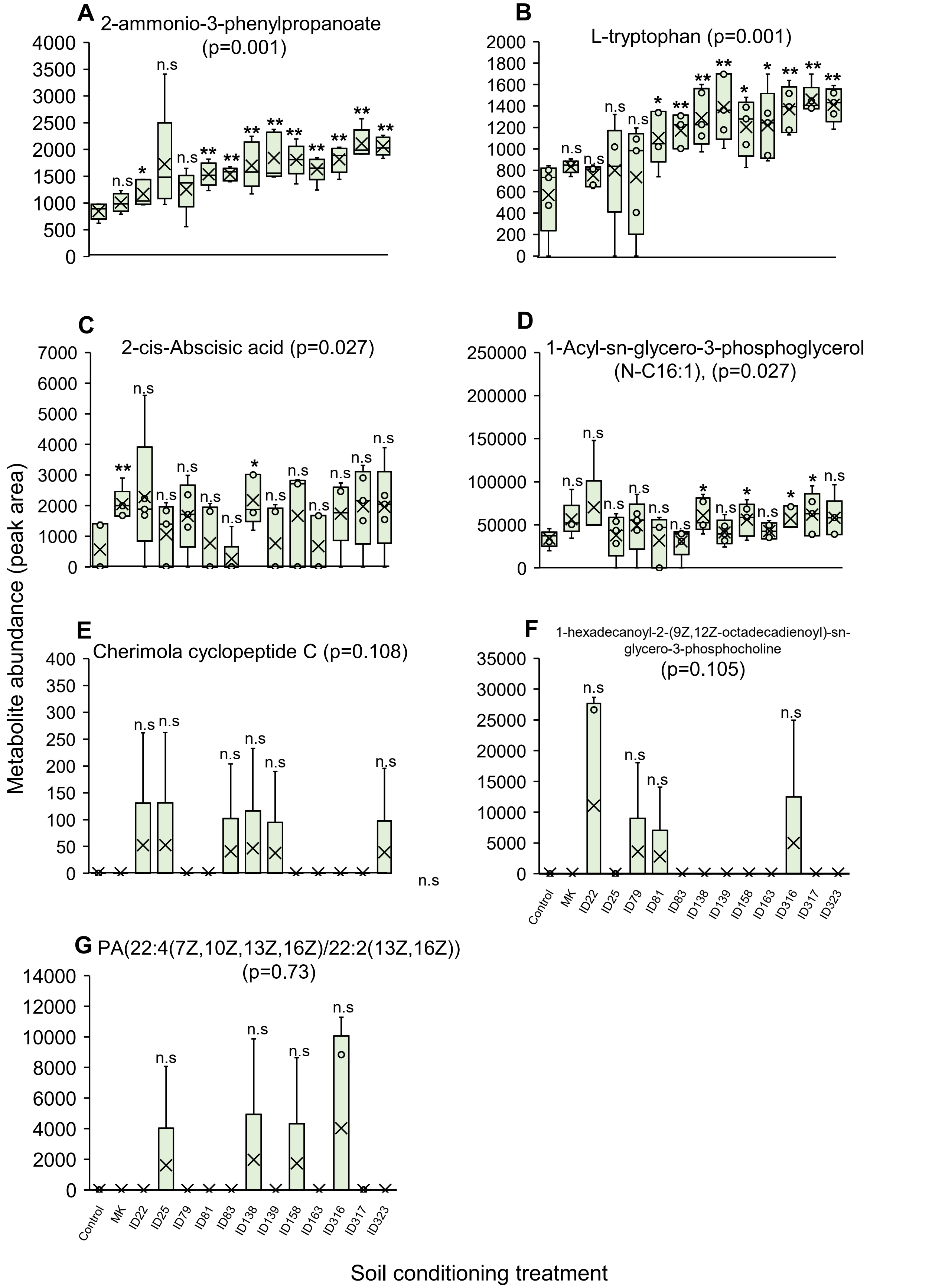


**Figure S21.** Plant leaves respond at the metabolic level to soil conditioning treatments. (**A–G**) Abundance of upregulated low molecular weight metabolites in roots of plants grown on non-conditioned control soil and in soils conditioned with mechanically killed larvae (MK) or with *Photorhabdus*-infected insect cadavers. IDs 22–323 refer to the different *Photorhabdus* strains used for larval infections (Table 1). Boxes represent the interquartile range (25th – 75th percentiles). In the boxplots, dots represent the raw data, black crosses (X) indicate the median values of all variables, and whiskers indicate the minimum and maximum values. Asterisks above the bars indicate significant differences relative to the control ($p$ < 0.05), assessed by one-way ANOVA followed by Tukey's HSD test. n.s. not statistically significant.

**Table S1.** List of metabolites that are upregulated in response to soil conditioning treatments in plants that resist the attack of *D. balteata* larvae (Roots) or the attack of *S. frugiperda* larvae (leaves).

| ID | Metabolite | Family | m/z | p-value | Plant part accumulated in |
|---|---|---|---|---|---|
| 106 | 4-[(6R)-6-hydroxy-5,5-dimethylcyclohexen-1-yl]benzoic acid | Terpenoids | 245.1180082 | 0.036 | Roots |
| 130 | 2 cis-Abscisic Acid | Terpenoids | 265.1443499 | 0.04 | Roots |
| 152 | Aromadendrin | Alkaloids | 287.0558557 | 0.002 | Roots |
| 196 | Compound id 196 | Alkaloids | 305.1502976 | 0.008 | Roots |
| 501 | Imidazole alkaloids | Alkaloids | 413.1447281 | 0.003 | Roots |
| 580 | Canthoside B | Unknown | 445.134756 | 0.002 | Roots |
| 285 | (6R,9R,10R)-6,9,10-trihydroxyoctadec-7-enoic acid | Fatty acids | 329.2332015 | 0.082 | Roots |
| 342 | Apigenin 7-sulfate | Shikimates and Phenylpropanoids | 349.0018846 | 0.057 | Roots |
| 953 | Pimarane and Isopimarane diterpenoids | Terpenoids | 655.3330362 | 0.14 | Roots |
| 978 | Ajugaciliatin A | Terpenoids | 669.3121824 | 0.304 | Roots |
| 334 | GDIMBOA | Benzoxazinoid | 344.0980292 | 0.487 | Roots |
| 891 | Tinosineside B | Alkaloids | 597.2182435 | 0.182 | Roots |
| 28 | 2-ammonio-3-phenylpropanoate | Amino acids and Peptides | 164.071264 | 0.001 | Leaves |
| 60 | L-Tryptophan | Amino acids and Peptides | 203.0823064 | 0.001 | Leaves |
| 663 | 1-Acyl-sn-glycero-3-phosphoglycerol (N-C16:1) | Fatty Acids | 481.2571387 | 0.027 | Leaves |
| 785 | 2 cis-Abscisic Acid | Terpenoids | 549.2441131 | 0.027 | Leaves |
| 1003 | Cherimola cyclopeptide | Amino acids and Peptides | 691.3543569 | 0.108 | Leaves |
| 1113 | PC(16:0/18:2(9Z,12Z)) $ 1-hexadecanoyl-2-(9Z,12Z-octadecadienoyl)-sn-glycero-3-phosphocholine | Shikimates and Phenylpropanoids | 802.5595557 | 0.105 | Leaves |
| 1115 | PA(22:4(7Z,10Z,13Z,16Z)/22:2(13Z,16Z)) | Unknown | 803.56318 | 0.73 | Leaves |
